# Extracellular matrix and cytoskeletal reverse remodeling pathways are key drivers of myocardial recovery following left ventricular assist device therapy

**DOI:** 10.1101/2025.11.06.687098

**Authors:** Thirupura S. Shankar, Marco Marchetti, Harini Srinivasan, Joseph R. Visker, Rana Hamouche, Ezra Johnson, James Jeong, Ioannis Kyriakoulis, Jing Ling, Konstantinos Sideris, Craig H. Selzman, Edgar J. Hernandez, Martin Tristani-Firouzi, Eleni Tseliou, Stavros G. Drakos, Omar Wever-Pinzon

**Affiliations:** Nora Eccles Harrison Cardiovascular Research and Training Institute, University of Utah, Salt Lake City, Utah; Eccles Institute of Human Genetics, University of Utah, Salt Lake City, Utah; Division of Cardiovascular Medicine, University of Utah, Salt Lake City, Utah; Division of Cardiothoracic Surgery, University of Utah, Salt Lake City, Utah; Division of Pediatric Cardiology, University of Utah, Salt Lake City, Utah; Division of Cardiology, University of Miami Health, Miami, Florida; Jackson Memorial Hospital, Miami, Florida; Miami Transplant Institute, Miami, Florida

**Keywords:** Whole genome bisulphite sequencing, RNA sequencing, epigenetics, Responder, Non-responder, Cardiac recovery, Heart failure

## Abstract

Transcriptomic changes in heart failure (HF) patients prior to and following left ventricular assist device (LVAD) support have been extensively studied. Recent studies focused on understanding DNA methylation changes in patients with cardiovascular diseases (CVD) and the role of circulating markers of DNA methylation as clinical predictors of the risk of CVD related morbidity and mortality. In this study, we used paired (pre- and post-LVAD) myocardial samples to examine changes in DNA methylation alongside RNA and protein expression. Our data suggests that patients with no improvement in cardiac function after LVAD therapy, despite showing an improvement in energy production (increased β-oxidation of fatty acid) exhibited persistent activation of profibrotic signaling, increased collagen deposition and cytoskeletal disarray evident from abnormal increase in sarcomeric distance following LVAD support. Contrarily, patients with improvement in cardiac function after LVAD therapy showed activation of pro-inflammatory signaling, collagen degradation and myogenesis. Both RNA sequencing and western blot data showed increased COL1A1 and decreased TPPP3 in post-NR thereby suggesting increased fibrosis and disrupted cytoskeletal signaling as potential barriers to myocardial recovery. Additionally, responders to LVAD therapy showed a significant reversal in myocardial interstitial fibrosis with a preserved sarcomeric architecture. Mice model of HF and recovery also confirmed our human findings, with reduced fibrotic signaling and improved cytoskeletal remodeling signaling observed in mice that showed improvement in cardiac function compared to mice with HF. Overall, our data suggests that altering extracellular matrix regulation and cytoskeletal signaling pathways may contribute to myocardial recovery. Further studies targeting these pathways are required to identify new HF therapeutic targets.

## Introduction

Heart failure (HF) is a global epidemic that affects over 56.5 million people,^1^ leading to significant morbidity and expense to healthcare systems. The prognosis of HF patients remains poor, with a 5-year survival rate of 50% and a 10-year survival rate of only 30%^2^. HF is a clinical syndrome with significant heterogeneity that includes diverse patient phenotypes with different underlying mechanisms and molecular traits, leading to varied clinical courses and different responses to therapy. Such heterogeneity poses challenges to the standard ‘one-size fits all’ approach, which typically involves initiation of guideline directed medical and device therapy (GDMT)^3^, followed by the use of advanced therapies such as left ventricular assist devices (LVADs) and transplantation in patients that fail to respond to GDMT^3^. To address this challenge, personalized strategies involving ‘omics’ and data-driven deep phenotyping could aid in the development of targeted therapeutics. Studies in the past decade have shown that a subset of advanced HF patients on LVAD support exhibit substantial improvement in both cardiac structure and function to allow for device removal (i.e. cardiac recovery and remission from chronic HF), herein classified as responders (R) (approximately 10-15% of advanced HF therapy candidates), whereas the remaining majority exhibits a continuous spectrum of changes ranging from moderate levels of improvement (i.e. partial responders, approximately 30% of advanced therapies candidates) to no improvement or continue to deteriorate and are classified as non-responders (NR, approximately 50% of advanced therapies candidates) ^4,5,6,7^. Intensive research has been ongoing to elucidate the mechanisms implicated in these favorable functional and structural changes that could lead to new HF therapies^8,9,10^.

Transcriptomic changes following LVAD unloading have been well documented, including a downregulation of fetal cardiac reprogramming and extracellular matrix (ECM) remodeling, along with an upregulation of oxidative phosphorylation and energy metabolism^11,12^. These transcriptomic changes are mediated by underlying epigenetic modifications, including DNA methylation. Changes in DNA methylation have been leveraged as a predictive tool, allowing the development of a blood-based risk score that stratifies individuals by their likelihood of developing CVD^13,14^. It has also been used to elucidate the pathogenesis of HF, including work from our group in which we identified changes in DNA methylation (and corresponding changes in gene expression) of genes involved in metabolic remodeling in HF, including hypermethylation of NRF1, which contributes to oxidative metabolic gene suppression in HF metabolism^15^. Likewise, other studies have identified DNA methylation changes in key genetic pathways in failing vs non-failing hearts^16,17,18^. Taken together, these data support an important role for DNA methylation patterns as an upstream regulatory mechanism and as a potential target for therapy development. Despite substantial research on DNA methylation changes in HF, the role of these epigenetic changes in myocardial recovery remains poorly explored^19^.

In the present study, we performed paired whole-genome bisulphite sequencing (WGBS) and bulk RNA sequencing in HF patients prior to and after a period on LVAD support. Patients were classified as responders and non-responders based on their myocardial structural and functional response to LVAD therapy^4^. Our data suggests that genes involved in tubulin/microtubule signaling and ECM reverse remodeling are implicated in the functional and structural myocardial recovery observed in responders following LVAD support. Furthermore, in responders to LVAD therapy we observed enrichment in genes related to improved cardiac metabolism and inflammatory response, including enhanced fatty acid oxidation and cardioprotective interleukin signaling. To validate these finding, we used a mouse model of HF (transverse aortic constriction, TAC) and recovery (de-TAC), which recapitulated the transcriptomic features of human LVAD responders.

Overall, our data showed that altering extracellular matrix regulation and cytoskeletal signaling pathways may contribute to myocardial recovery. Future studies targeting these pathways may provide more insights into the molecular mechanisms underlying cardiac recovery and pave way for novel therapeutic interventions.

## Methods

### Study population

Patients with advanced HF were prospectively enrolled at the time of LVAD implantation. This study was carried out at institutions comprising the Utah Cardiac Recovery Program (UCAR) and the Utah Transplant Affiliated Hospitals (U.T.A.H) Cardiac Transplant Program (University of Utah Health, Intermountain Medical Center, and the Salt Lake VA Medical Center, all in Salt Lake City). The institutional review board of each institution approved the study, and all patients provided informed consent. Baseline characteristics are included in Table 1 where the continuous and ordinal variables were summarized as median [IQR] and compared between responders and non-responders using the Wilcoxon rank-sum test. Categorical variables were summarized as n (%) and compared using Fisher’s exact test for 2×2 tables or Pearson’s χ² test for multi-level categories. All tests were two-sided (α=0.05) and performed in Stata 16. As previously known, we observed a significant difference in the age of the population, with responders being younger than non-responders^20,21^. We also observed a significant difference in VAD indication between the groups, with more responders in the “bridge-to-recovery” group as predicted. No significant difference in etiology, medications, LVAD type, etc. was observed between the two groups.

**Table 1:** Patient characteristics.

|  | **Non-responders**<br>**(n=56)** | **Responders**<br>**(n=11)** | **p-value** |
| --- | --- | --- | --- |
| ***Demographics*** |  |  |  |
| Age (years) | 56.0 [42.5–60.5] | 38.0 [19.0–60.0] | **0.027** |
| BMI (kg/m<sup>2</sup>) | 27.29 [24.28–31.47] | 28.03 [22.18–34.43] | 0.732 |
| ***Echocardiography*** |  |  |  |
| LVEF (%) pre | 15.0 [10.0–20.0] | 15.0 [12.0–20.0] | 0.923 |
| LVEF (%) post | 25.0 [20.0–28.0] | 45.0 [40.0–48.0] | **<0.001** |
| LVEDD (cm) pre | 6.77 [6.26–7.49] | 6.81 [5.80–7.20] | 0.573 |
| LVEDD (cm) post | 6.04 [5.21–6.80] | 4.83 [4.74–5.70] | **0.003** |
| ***HF history / severity*** |  |  |  |
| HF duration (months) | 72.0 [13.5–120.0] | 22.0 [4.5–120.0] | 0.132 |
| NYHA class (pre-LVAD) |  |  | 0.628 |
| III | 14/56 (25%) | 2/11 (18.2%) |  |
| IV | 42/56 (75%) | 9/11 (81.8%) |  |
| **INTERMACS profile, median [IQR]** | **3 [3–4] (n=55)** | **3 [2–4] (n=11)** | **0.22** |
| Pre-implant CHF = Acute (Yes) | 8/56 (14.3%) | 3/11 (27.3%) | 0.371 |
| Inotropes dependent | 40/55 (72.7%) | 8/11 (72.7%) | 1.000 |
| ***Etiology*** |  |  | 0.304 |
| Ischemic etiology (ICM) | 22/56 (39.3%) | 2/11 (18.2%) | 0.304 |
| Non ICM | 34/56 (61.7%) | 9/11 (81.8%) |  |
| ***Comorbidities*** |  |  |  |
| Smoking history | 32/56 (57.1%) | 5/11 (45.5%) | 0.523 |
| ETOH history | 30/56 (53.6%) | 4/11 (36.4%) | 0.340 |
| Diabetes | 16/56 (28.6%) | 1/11 (9.1%) | 0.266 |
| Hypertension | 24/56 (42.9%) | 5/11 (45.5%) | 1.000 |
| Atrial fibrillation | 21/56 (37.5%) | 1/11 (9.1%) | 0.086 |
| ***VAD type*** |  |  | 0.572 |
| Jarvik | 3/56 (5.4%) | 1/11 (9.1%) |  |
| HeartMate II | 16/56 (28.6%) | 3/11 (27.3%) |  |
| HeartMate III | 1/56 (1.8%) | 1/11 (9.1%) |  |
| HVAD | 36/56 (64.3%) | 6/11 (54.6%) |  |
| **Device therapy** |  |  | 0.691 |
| None | 8/56 (14.3%) | 2/11 (18.2%) |  |
| ICD | 25/56 (44.6%) | 6/11 (54.6%) |  |
| CRT-D | 23/56 (41.1%) | 3/11 (27.3%) |  |
| **VAD indication** |  |  | **<0.001** |
| Bridge to Transplant | 54/56 (96.4%) | 4/11 (36.4%) |  |
| Bridge to Recovery | 0/56 (0.0%) | 5/11 (45.5%) |  |
| Bridge to Decision | 2/56 (3.6%) | 2/11 (18.2%) |  |
| **Medications post-LVAD** |  |  |  |
| Beta blocker | 34/56 (60.7%) | 10/11 (90.9%) | 0.082 |
| ACEi/ARB | 38/56 (67.9%) | 8/11 (72.7%) | 1.000 |
| Aldosterone blocker | 34/56 (60.7%) | 7/11 (63.6%) | 1.000 |
| Warfarin | 56/56 (100.0%) | 10/11 (90.9%) | 0.164 |
| Diuretics | 51/56 (91.1%) | 11/11 (100%) | 0.582 |

### Patient sample acquisition

LV myocardial samples were prospectively collected from the apical core (at the time of LVAD implantation) and close to the apex (at the time of transplant/ explant) as previously described and were snap frozen in liquid nitrogen^22^. The WGBS and RNA sequencing data reported in our study were generated from male samples. Preliminary analyses revealed clear sex-based differences, however, due to limited number of female samples, we focused on male data to ensure statistical robustness. We plan to perform a follow up study dedicated to investigating these sex-specific differences. In contrast, the western blots, Masson’s trichrome and TEM analyses includes both male and female due to the small sample size available for these experiments.

### Animals and animal care

All animal studies were performed in accordance with the Institutional Animal Care and Use Committee (IACUC). All procedures involving animals were approved by the Animal Care and Use Committee of the University of Utah and complied with the American Physiological Society’s *Guiding Principles in the Care and Use of Animals* and the UK Animals (Scientific Procedures) Act 1986 guidelines. The mice were housed in 12 h dark/light cycle at 70 °F and 40% humidity. Both male and female mice were used in this study.

### RNA extraction, sequencing and analysis

Total RNA was extracted from left ventricular (LV) myocardial samples from both mouse and human tissue, using the miRNeasy Mini Kit (Qiagen, REF 217084), with on-column DNase digestion performed using the RNase-Free DNase Kit (Qiagen, REF 79254) to eliminate genomic DNA. Library preparation was performed using the Illumina TruSeq Stranded RNA Kit with Ribo-Zero Gold for rRNA depletion and sequenced on an Illumina HiSeq 2500 platform using 50 bp single-end reads. Raw reads were trimmed using TrimGalore (0.6.10, https://github.com/FelixKrueger/TrimGalore) prior to alignment to the human (hg38, Ensembl 104) or mouse (m39, Ensembl 104) genome using STAR (2.7.11a)^23^ with the "--quantMode GeneCounts" option. Strong batch effects arising from samples being sequenced in different batches were observed, then corrected using the ComBat_seq function from the R package SVA (3.50.0)^24^ package. Differential expression analysis was performed using the R package DESeq2 (v1.42.1)^25^. In line with established practices in genomic biomarker studies, genes with a Benjamini-Hochberg corrected p value < 0.2 were considered as significantly differentially expressed and were using for downstream analyses using Ingenuity pathway analysis (IPA)^26,27^.

### Gene set score computation

A gene set score represents the aggregate expression of a set of genes of interest in a sample compared to the expression of a control set. The control set is identified as a random subset of genes that has similar expression to the genes of interest when averaged across all samples. The gene set score indicates whether a gene set is expressed in a specific sample more or less than in the average of the samples. First, gene expression counts for each sample were normalized for total sample counts, log1p-transformed, then normalized again for transformed total counts. Counts for each individual gene were then Z-score-normalized across samples. Gene set scores were computed for each sample using these transformed and scaled counts as follows: 1) gene expression was averaged across samples and binned in a user-defined number of bins; 2) a control set was identified as a set of randomly selected genes that in the whole sample population falls in the same bins as genes from the set of interest was identified; 3) the expression of the genes in the control set in the sample of interest was averaged; 4) the expression of the genes in the target set in the sample of interest was also averaged; 5) the control set averaged expression was subtracted from the target gene set averaged expression. The value obtained is the gene set score for the specific sample. Gene set scores were then compared across conditions by Mann-Whitney U test.

### WGBS sequencing and analysis

TrimGalore (0.6.10) was used to pre-process raw reads by trimming adapters and removing low-quality (PHRED <20) reads. Bismark (0.24.0)^28^ was then used for alignment to the human genome (hg38, Ensembl 104), deduplication, and methylation call extraction. To quantify differential DNA methylation of CpG islands, the R package methylKit (1.28.0)^29^ was used. Briefly, methylation calls were merged at the level of know CpG islands, filtered for too low (< 10x) or high (> 99.9 percentile) depth, normalized, united across samples, corrected for batch-effects with the assocComp function from methylKit, and finally a differential methylation analysis was run using a chi-square test with basic overdispersion correction. CpG islands were defined as significantly differentially methylated with q-value < 0.2. CpG islands were then annotated with overlapping genes within 2Kbp of their extremities.

### Protein extraction and Western blotting

30μg of human myocardial tissue and mouse cardiac tissues were used for this experiment. Protein extraction and western blotting were performed as previously described.^30,31^ Briefly, tissues were homogenized in RIPA buffer containing 2× protease and phosphatase inhibitor (ThermoScientific #1861281) using metal beads for 3 minutes. The homogenate was transferred to a fresh tube, with 10μl of 1× PMSF (0.1M), and kept for rotation at 4°C for 30 minutes. Samples were then centrifuged at 11,000×g for 10 minutes at 4°C, and the supernatant was collected for protein quantification using the Pierce BCA Protein Assay Kit (ThermoScientific #23225) Protein lysates were diluted with equal volume of 2× Laemmli buffer containing 10% DTT, followed by boiling at 98°C for 10 minutes.

Proteins (30μg per sample) were separated via SDS-PAGE at a constant voltage of 25V/gel and transferred to nitrocellulose membranes at 350mA. Membranes were blocked for 1 hour in 5% non-fat milk and incubated with primary antibodies (Rabbit anti-Collagen1a1 (Novus biologicals #NB600-408, 1:100); Rabbin anti-TPPP3 (Proteintech #15057-1-AP, 1:100)) overnight at 4°C. The blots were then washed three times with 1× TBST (10 minutes each) and incubated with the appropriate secondary antibody (Donkey anti-Rabbit; Invitrogen #926-32213, 1:10000 in 1x TBST) for 1 hour. Membranes were then washed again three times with 1× TBST (10 minutes each). Blots were scanned using the LI-COR imaging system, and Image Studio Lite software v5.2 was used for protein quantification. Each blot included a lane-specific loading control, and protein expression values were normalized accordingly.

### Transmission Electron Microscopy (TEM)

Methodology for TEM sample preparation, staining, and analysis outlined are similar to our previous publication^30^. Left ventricular (LV) transmural tissue samples from R and NR groups were analyzed. Tissues were fixed overnight at 4 °C in 0.1 M sodium cacodylate buffer containing 1% paraformaldehyde and 2.5% glutaraldehyde. After fixation, samples were washed in the same buffer and post-fixed for 2 hours in 2% osmium tetroxide prepared in cacodylate buffer. Samples were rinsed with nano-pure water and stained with en-bloc with uranyl acetate for 1 hour at room temperature.

Dehydration was performed using a graded ethanol series as follows: 50% ethanol for 10 minutes, followed by three washes of 70% ethanol (10 minutes each), two washes of 95% ethanol (10 minutes each), four washes of 100% ethanol (10 minutes each), and three washes of absolute acetone (10 minutes each). The tissues were infiltrated with epoxy resin (Electron Microscopy Sciences, Hatfield, PA) in a stepwise manner. Samples were incubated at room temperature in 50% resin in acetone for 1 hour, then in 75% resin in acetone overnight, and finally in 100% resin for 8 hours with three changes of fresh resin.

Following infiltration, the tissues were embedded in resin and polymerized at 60 °C for 48 hours. Ultrathin sections (70 nm) were prepared using a diamond knife (Diatome) on a Leica UC6 ultramicrotome (Leica Microsystems, Vienna, Austria). The sections were post-stained with saturated uranyl acetate for 10 minutes, followed by 5 minutes in Reynold’s stain. Samples were imaged on a JEOL 1400 Plus transmission electron microscope. Sarcomere length was measured from the acquired images using Fiji software.

### Transverse Aortic Constriction (TAC) and de-TAC

12-week-old C57BL6J mice were used for this study. Mice were anesthetized using isoflurane placed on a heated pad. Depilatory cream was used to remove fur from the surgery area. Betadine and 70% EtOH was used to sterilize the surgical site and appropriate medications given as per IACUC protocol. Skin and muscle were cut to expose the trachea and a 6-0 prolene suture was used to constrict the aorta. We used a 33-gauge needle to gauge the level of constriction. Sham mice underwent the same procedure, but the aorta was not constricted. Serial echocardiography was performed on all mice and deTAC was performed on a random subset of mice that developed HF (EF<35%) to subsequently observe for evidence of cardiac recovery. For the deTAC mice, all pre-surgical procedures were the same as above, except, a fine-end scissor was used to remove the suture around the aorta. Same steps were repeated on the sham mice. Serial echocardiography was performed on the mice for 4-weeks and tissue harvested. Sham mice were not included in the data analysis since the comparisons were between mice with HF and mice that showed improved cardiac function following de-TAC.

### Echocardiography

Echocardiographic studies in humans were performed at the echocardiography laboratories of the participating institutions using a previously described protocol^32^ and using previously described techniques in accordance with current American Society of Echocardiography guidelines.

In mice, echocardiography was performed as previously discussed^33^. Briefly, mice were anesthetized using 1.5% isoflurane (Vet One, NDC 13985-046-60). Imaging was performed using the Vevo 2100 system. Two-dimensional (2D) short-axis views were acquired and analyzed using Vivo strain software (version 3.1.1). For accuracy and consistency, measurements were derived from two consecutive cardiac cycles. Additionally, electrocardiograms (ECG) were recorded via limb leads during all echocardiographic procedures.

### Mason’s Trichrome Staining

Paraffin blocks were used 8μm sections were cut. Staining was performed as per manufacturers protocol (Sigma-Aldrich Trichrome Stain (Masson) #HT15-1KT). Slides were scanned at ×20 magnification under a light microscope and analyzed using MATLAB_R2025b. The ratio of collagen-stained area to the total stained area was calculated and reported^34^.

## Results

### Reduced fibrotic signaling and improved cytoskeleton re-organization signaling observed following LVAD therapy

A total of 67 HF patients were included in this study (Table 1), with paired comparisons performed at pre- and post-LVAD timepoints (Figure 1A). RNA sequencing data revealed a total of 3323 genes significantly upregulated, and 3393 genes downregulated post-LVAD (Figure 1B). Amongst the most differentially expressed were ABRA, PEAK3, KLF15, PEAK3 and MSTN (upregulated) and COL1A1, RUN1, ELN (downregulated), suggesting shifts in cytoskeletal and fibrotic signaling. Ingenuity pathway analysis (IPA) of RNA sequencing data in all comers at post-LVAD timepoint compared to pre-LVAD revealed activation of pathways associated with cardiac remodeling, including apelin signaling in cardiac fibroblasts, VEGF pathway activation^35^, collagen trimerization, HIPPO and Notch signaling^36^, myogenesis and inhibition of matrix metalloproteases (MMPs). In contrast, pro-fibrotic and structural adhesion pathways including PI3K/AKT, integrin and adherens junction signaling were downregulated post-LVAD (Figure 1C-D). RNA sequencing data showed significant metabolic and inflammatory reprogramming post-LVAD evident from upregulation of fatty acid metabolism and downregulation of ceramide and glucose metabolism (Figure S1A-B). Patients also exhibited upregulation of cardioprotective IL13^37^ and IL22^38^ signaling following LVAD support, (Figure S1C-D). WGBS data identified 53 hypomethylated and 14 hypermethylated regions in post-LVAD samples compared to pre-LVAD (Figure S1E). Reactome pathway analysis of differentially methylated regions identified alterations in ECM reorganization and cytoskeletal signaling pathways. Specifically, in post-LVAD samples, we observed hypermethylation of genes involved in integrin signaling and hypomethylation of genes involved in cardioprotective pre-NOTCH signaling^39^ amongst other pathways (Figure 1E-F). Integrated analysis of RNA sequencing and WGBS data showed that genes exhibiting consistent patterns (e.g., hypomethylation in WGBS and simultaneous upregulation in RNA sequencing) were implicated in ECM remodeling and cytoskeleton signaling, highlighting these 2 pathways as key regulators of LVAD-mediated response (Figure 1G). Histological assessment of these tissues using Masson’s trichrome staining showed a significant reduction in myocardial interstitial fibrosis post-LVAD, confirming the impact of our molecular findings (Figure 1H). Transmission electron microscopy (TEM) on transmural myocardial samples from these patients showed no significant difference in the sarcomeric distance (measured as distance between the Z-lines) between pre- and post-LVAD samples suggesting the preservation of cytoskeletal architecture post-LVAD (Figure 1I). In summary, our data showed a substantial reduction in interstitial fibrosis and enhanced cytoskeleton signaling following LVAD therapy.

**Figure 1.**
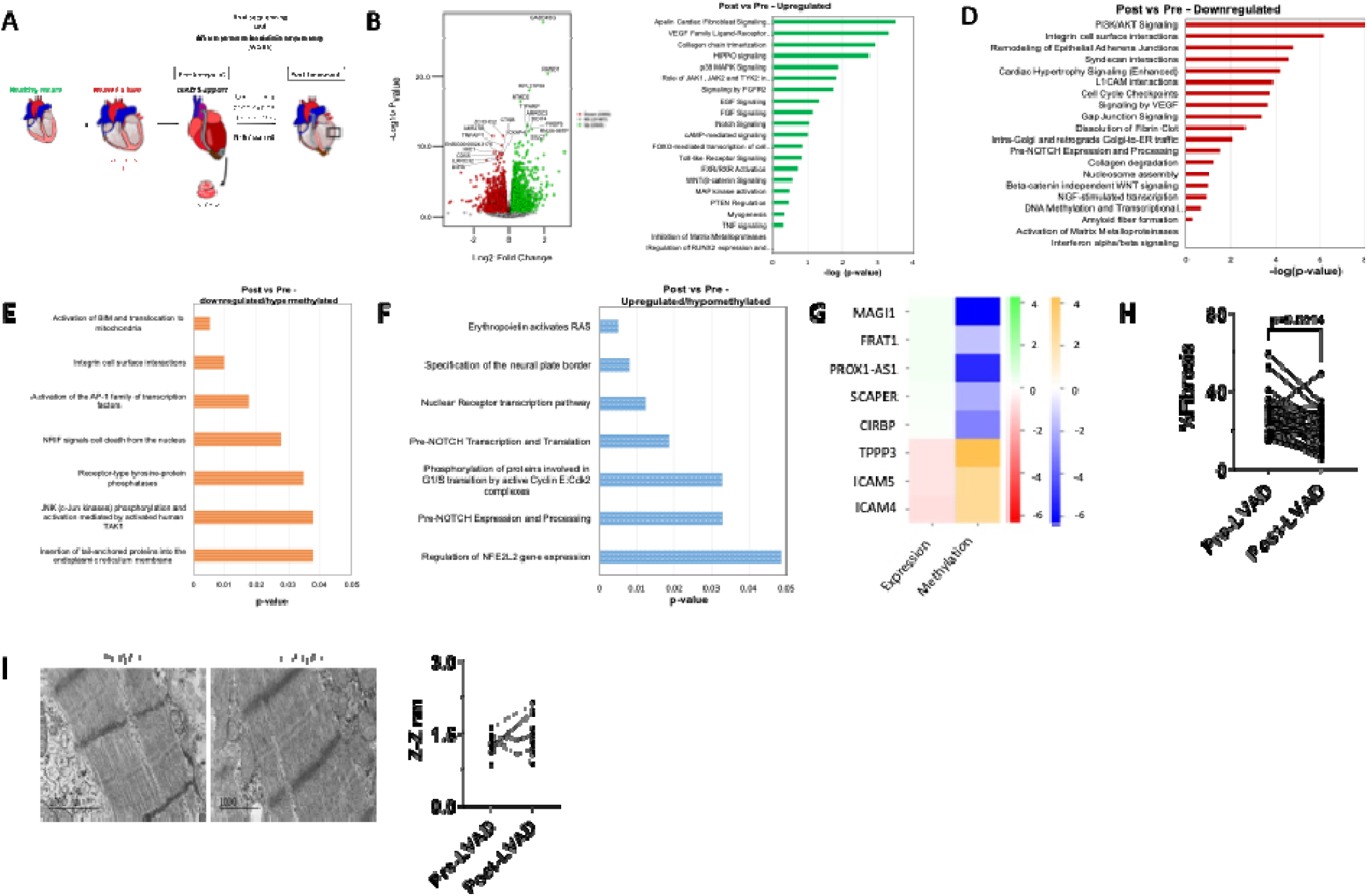
**A** Schematic of workflow for post-LVAD vs pre-LVAD comparisons (n=67 paired), **B.** Volcano plot of RNA sequencing data showing differentially expressed genes in post-LVAD vs pre-LVAD, **C-D.** Ingenuity pathway analysis of RNA sequencing data showing enriched pathways among upregulated and downregulated genes, **E-F.** Reactome pathway analysis showing enriched pathways among hypermethylated and hypomethylated genomic regions, **G.** Heatmap of RNA sequencing and WGBS data showing genes with concordant changes, where increased expression is associated with hypomethylation and decreased expression with hypermethylation, **H.** Masson’s trichrome quantification (n=34 paired), **I.** Representative transmission electron microscopy image and quantification of distance between the Z-lines (n=12). p-value: paired t-test.

### Increased fibrotic signaling, ECM remodeling and impaired actin cytoskeleton remodeling observed in non-responders (NR) following LVAD therapy

Patients whose LVEF remained unchanged or reduced were designated as NR (n=56 paired) (Figure 2A). RNA sequencing data showed a total of 2961 significantly upregulated, and 3421 genes downregulated in post-NR compared to paired pre-NR (Figure 2B). Amongst the most differentially expressed genes were DDIT4, RASD1, FKBP5 (upregulated) and LDHA, SMAD7 (downregulated) (Figure 2B). IPA revealed activation of pathways related to senescence, fibrotic signaling and apoptosis, while downregulation of ECM organization, gap junction and actin cytoskeleton signaling were observed in post-NR compared to their pre-LVAD timepoint (Figure 2C-D). Increased fatty acid oxidation and reduced glycolysis was observed in post-NR. The classical pentose phosphate pathway and sphingolipid signaling, which have been shown to be crucial for normal cardiac functioning^40,41^, were downregulated in post-NR compared to pre-NR (Figure S2A-B). Anti-inflammatory cytokine signaling was also upregulated in post-NR (Figure S2C-D). WGBS analysis revealed hypermethylation of 9 regions and hypomethylation of 2 regions in post-NR (Figure S2E). Reactome pathway analysis of differentially methylated regions revealed enrichment of pathways related to suppression of apoptosis and activation of fetal cardiac development, such as the formation of intermediate mesoderm and neural plate border, in post-NR (Figure 2E-F). In evaluating concordance between transcript expression and methylation, TPPP3 (tubulin polymerization promoting protein) was significantly hypermethylated and downregulated in post-NR, suggesting a reduction in microtubule bundling and consequently impaired cellular structure and mechanics^42,43^. Conversely, MAGI1 (a scaffolding protein at cell-cell junction) was hypomethylated and upregulated indicating increased apoptosis, endothelial activation and endoplasmic reticulum (ER) stress in post-NR^44^ (Figure 2G). Histological analysis showed no significant difference in absolute fibrosis levels between pre- and post-NR (Figure 2H). However, post-NR samples manifested a significant increase in sarcomeric length suggestive of stretched muscle fibers and impaired cardiac contraction compared to the pre-LVAD timepoint^45^ (Figure 2I). Overall, our data suggests that patients without improvement in their myocardial structure and function following LVAD therapy exhibited activation of pathways associated with increased fibrosis, ECM remodeling, and compromised actin cytoskeleton function.

**Figure 2.**
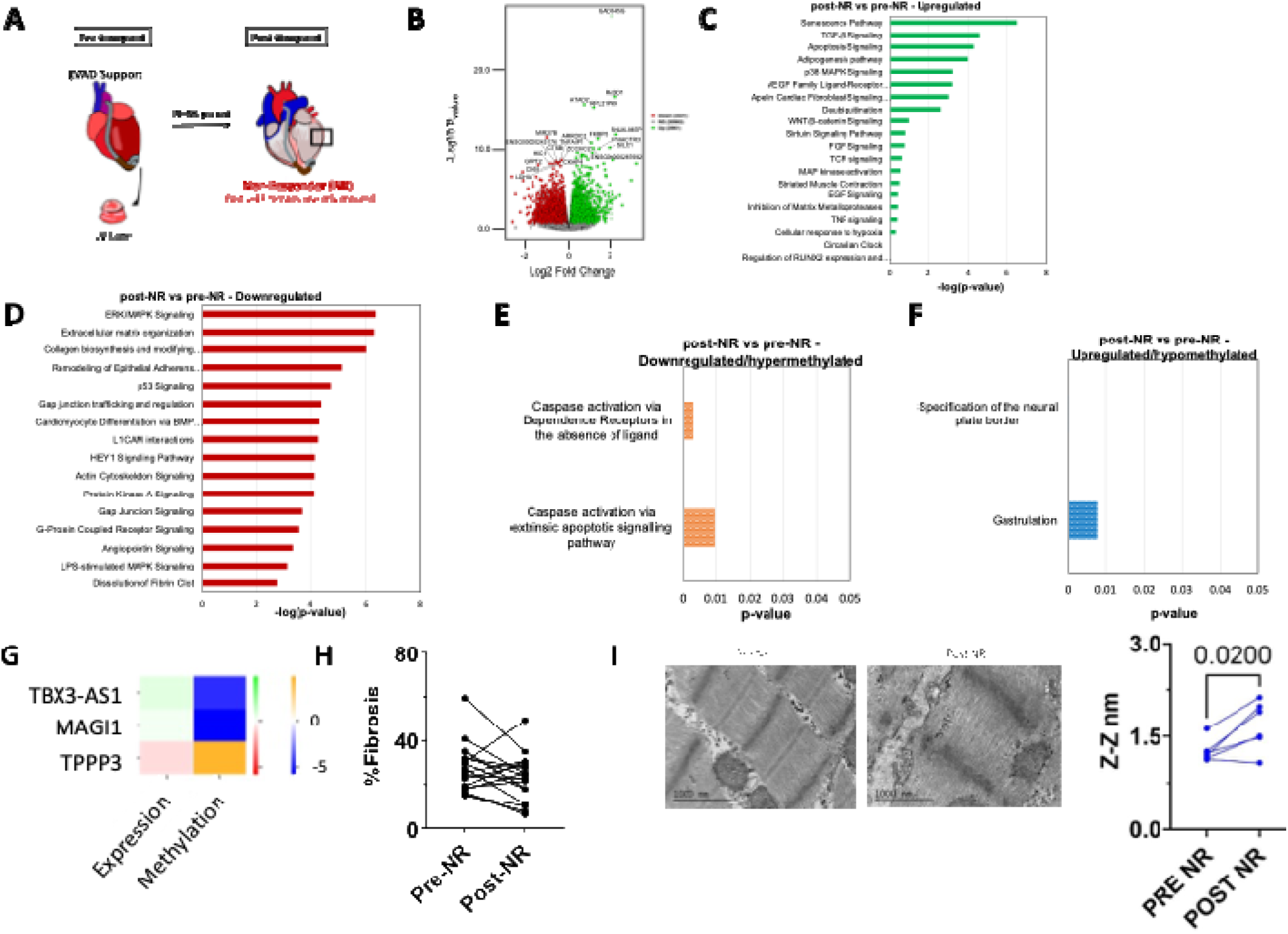
**A** Schematic of workflow for post-NR vs pre-NR comparison (n=56 paired), **B.** Volcano plot of RNA sequencing data showing differentially expressed genes between post-NR vs pre-NR (n=56 paired), **C-D.** Ingenuity pathway analysis of RNA sequencing data showing enriched pathways among upregulated and downregulated genes, **E-F.** Reactome pathway analysis showing enriched pathways among hypermethylated and hypomethylated genomic regions, **G.** Heatmap of RNA sequencing and WGBS data showing genes with concordant changes, where increased expression is associated with hypomethylation and decreased expression with hypermethylation, **H.** Masson’s trichrome quantification (n=17 paired), **I.** Representative transmission electron microscopy image and quantification of distance between the Z-lines (n=6). p-value: paired t-test.

### ECM reverse-remodeling and myogenesis pathways activated in responders post-LVAD therapy

Patients who exhibited both an improvement in cardiac structure and function were classified as responders (R) (Figure 3A). Differential expression analysis identified 842 upregulated and 730 downregulated genes in post-R compared to pre-R. Genes including SNAI2, PTPN2, DCN and MDK were significantly upregulated and TANGO2, MTOR and MAPK4 were significantly downregulated in post-R compared to their pre-LVAD timepoint (Figure 3B). IPA showed enrichment of pathways related to myogenesis, sirtuin signaling, collagen degradation, HIPPO signaling and wound healing in post-R, suggesting a classic ECM reverse-remodeling response. Furthermore, significant downregulation of apoptosis, pro-fibrotic pathways, integrin and NFAT signaling^46^ pathways were observed, suggesting a unique cardioprotective phenotype in responders following LVAD support. Interestingly, post-R showed a significant upregulation of pathways involving glycosaminoglycan metabolism and a significant downregulation of glycolysis and ceramide signaling following LVAD unloading (Figure S3A-B). Pro-inflammatory signaling pathways, IL-1^47^ and IL-15^48^, that protect the heart from adverse remodeling, were significantly upregulated in post-R, while a downregulation of IL-6 and IL-10, amongst other inflammatory mediators involved in adverse cardiac remodeling was observed in post-R compared to pre-R (Figure S3C-D). Analysis of WGBS data showed hypomethylation of 12 regions and hypermethylation of 9 regions in post-R compared to pre-R. A significant hypermethylation of KIF26A, TUBB3 and AGRN (genes involved in microtubule and laminin activity) and a significant hypomethylation of PKM and KL (genes involved in glycolysis and klotho signaling) were observed in post-R (Figure S3E). Activation of these genes in the heart has been shown to be cardioprotective^49,50^. Additionally, Reactome pathway analysis of differentially methylated regions revealed hypermethylation of genes in pathways associated with unstable tubulin folding intermediates and microtubule-dependent trafficking of connexons, along with hypomethylation of β-catenin destruction complex degradation and glucose metabolism gene pathways (Figure 3 E-F). Masson’s trichrome staining showed a significant reduction in myocardial interstitial fibrosis levels. No difference in the sarcomeric length was observed in the post-R compared to paired pre-R, indicating stable structural integrity of sarcomeres in responders (Figure 3G-H). In summary, our findings suggest that responders to LVAD therapy exhibit genomic and structural changes consistent with ECM reverse remodeling, enhanced cardioprotective signaling, and improved cardiac function.

**Figure 3.**
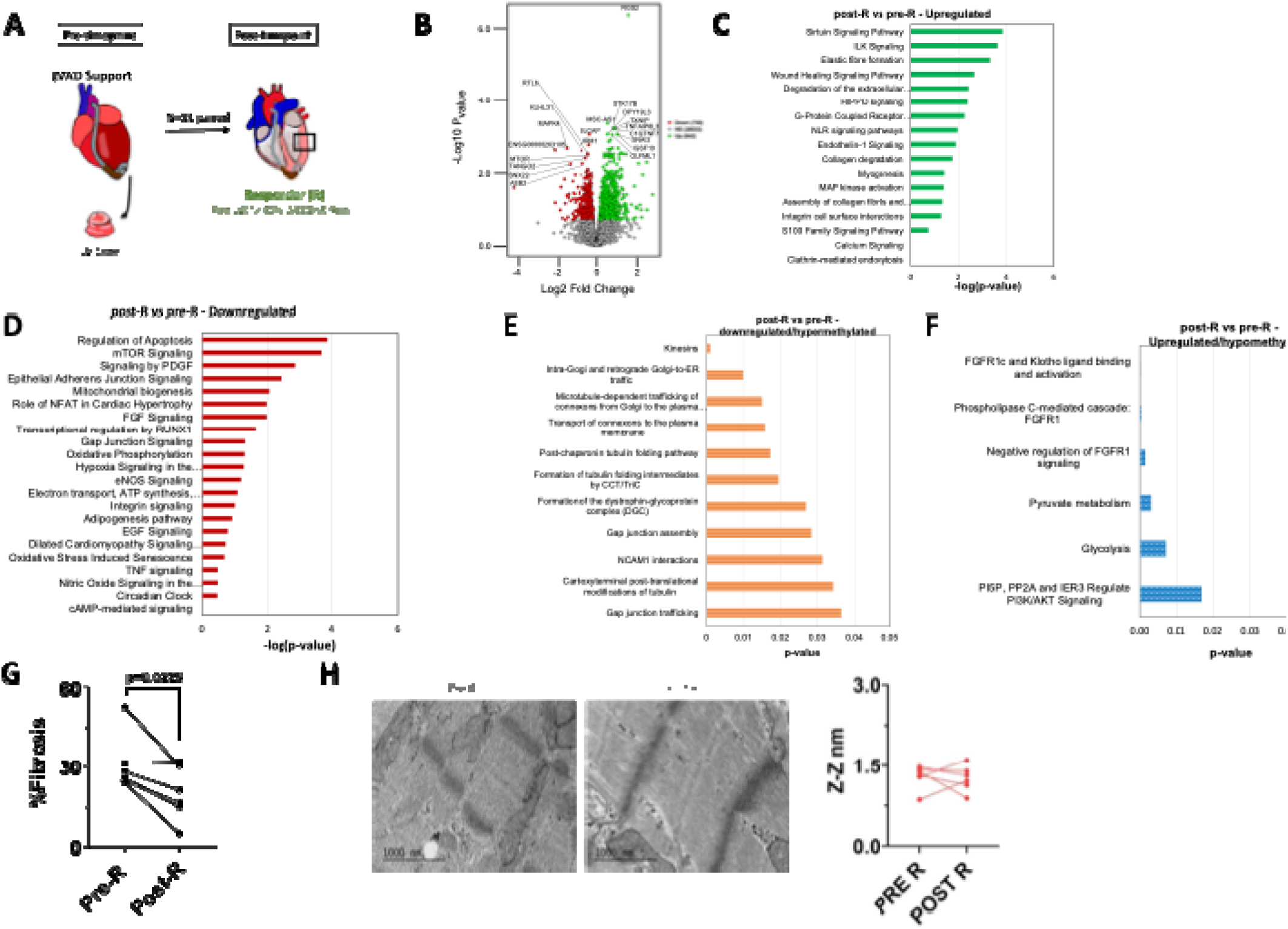
**A** Schematic of workflow for post-R vs pre-R comparison (n=11 paired), **B.** Volcano plot of RNA sequencing data showing differentially expressed genes between post-R vs pre-R (n=11 paired), **C-D.** Ingenuity pathway analysis of RNA sequencing data showing enriched pathways among upregulated and downregulated genes, **E-F.** Reactome pathway analysis showing enriched pathways among hypermethylated and hypomethylated genomic regions, **G.** Masson’s trichrome quantification (n=6 paired), **I.** Representative transmission electron microscopy image and quantification of distance between the Z-lines (n=6). p-value: paired t-test.

### Responders are potentially predisposed to reduced fibrotic signaling and improved beta-oxidation of fatty acids compared to non-responders prior to LVAD implantation

Despite receiving similar hemodynamic support with an LVAD, a subset of patients experience significant myocardial improvement, both structurally and functionally, while the majority show less improvement, stability or experience further deterioration (i.e. a continuous spectrum or responses). To determine whether responders are predisposed (or ‘primed’) to recover function following LVAD therapy, we compared gene expression profiles of pre-R and pre-NR (Figure 4A). Differential gene expression analysis identified 127 significantly downregulated genes and 5 upregulated genes. Notably, genes including GOLGA8S, KLHL1 and PKD1L1 were significantly upregulated while FOS, VCAM2, HAS2 and DUSP4 were significantly downregulated in pre-R compared to pre-NR (Figure 4B). PPAR signaling and anti-inflammatory signaling pathways were upregulated in pre-R, whereas pathways associated with fibrosis and hypertrophy were downregulated in pre-R relative to pre-NR (Figure 4C-D, Figure 4SA-D). WGBS data showed 133 hypomethylated and 594 hypermethylated regions in pre-R compared to pre-NR (Figure 4SE). Genes in pathways related to cell cycle and apoptosis-induced DNA fragmentation were hypermethylated, while genes in cardioprotective pathways, including FOXO-mediated signaling and AKT phosphorylation, were hypomethylated (Figure 4E-F). FOXO and AKT pathways were previously shown to be crucial for cardioprotective signaling and promoting cardiac cell survival following injury such as myocardial infarction (MI)^51,52^. In addition, genes involved in MAPK signaling (DUSP6, MAP3K8) and JUNB were hypermethylated and downregulated reinforcing the presence of a strong cardioprotective phenotype in responders prior to LVAD therapy (Figure 4G). No significant in fibrosis levels or sarcomere distance at baseline was observed between responders and non-responders, suggesting comparable structural derangement (Figure 4H). Overall, our data indicates that responders to LVAD exhibit distinct gene expression and methylation signatures prior to LVAD implant suggestive of an inherent predisposition to myocardial recovery.

**Figure 4.**
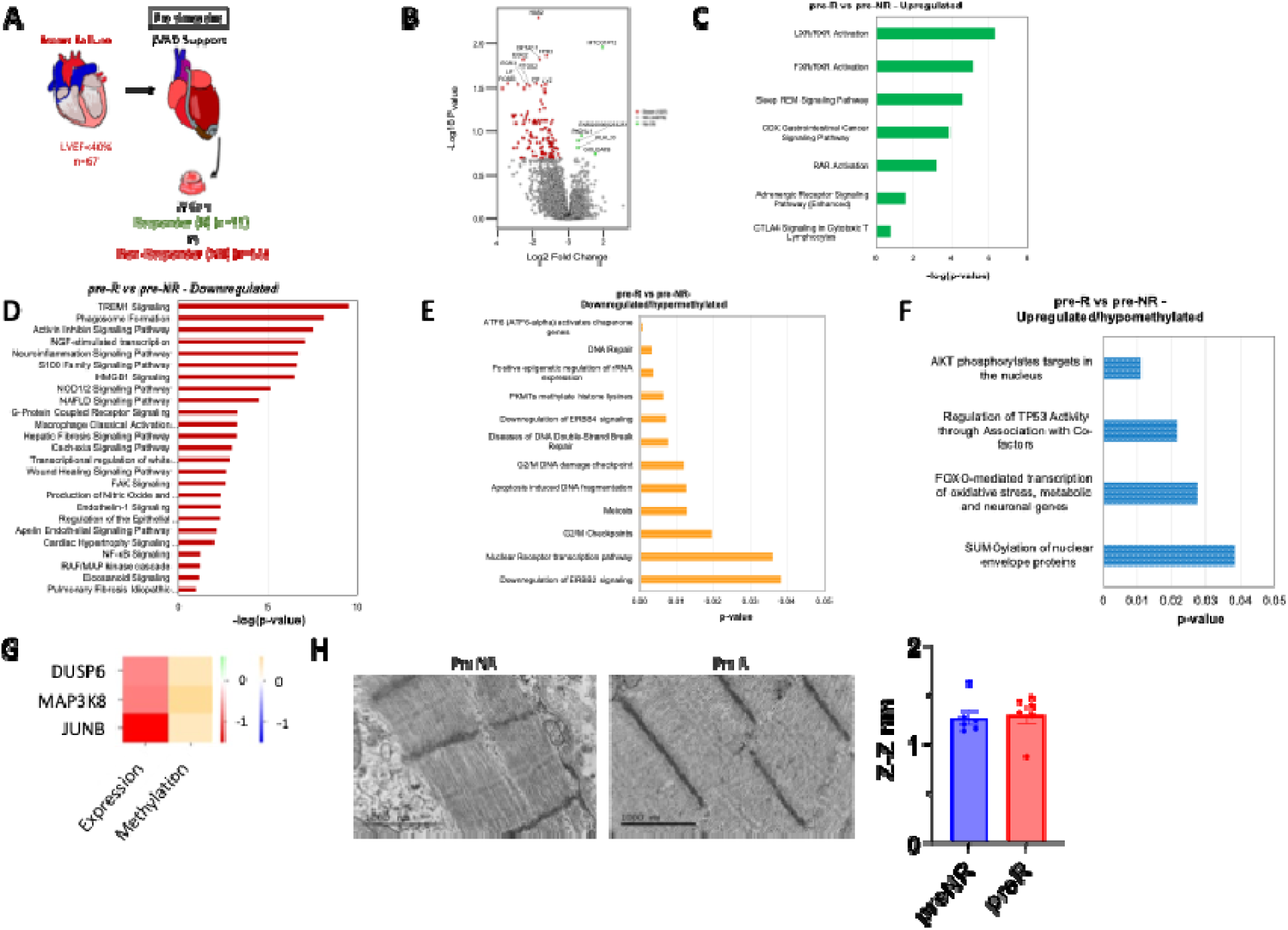
**A** Schematic of workflow for pre-R (n=11) vs pre-NR (n=56) comparison, **B.** Volcano plot of RNA sequencing data showing differentially expressed genes between pre-R vs pre-NR, **C-D.** Ingenuity pathway analysis of RNA sequencing data showing enriched pathways among upregulated and downregulated genes, **E-F.** Reactome pathway analysis showing enriched pathways among hypermethylated and hypomethylated genomic regions, **G.** Heatmap of RNA sequencing and WGBS data showing genes with concordant changes, where increased expression is associated with hypomethylation and decreased expression with hypermethylation, H. Representative transmission electron microscopy image and quantification of distance between the Z-lines (n=6). p-value: unpaired t-test.

### Reduced fibrosis and tubulin signaling potentially drives myocardial recovery following LVAD support

To further validate the differential responses to LVAD unloading, we compared post-R to post-NR, tracking changes in gene expression and methylation patterns (Figure 5A). Differential expression analysis revealed a significant upregulation of 28 genes and downregulation of 130 genes in post-R compared to post-NR. Amongst these, we observed a significant increase in the expression of wound-healing related genes (MMP2, LAMC1, DDR2) and a downregulation of contractility-related genes (TPM2, TPM1) in post-R (Figure 5B). Pathways related to dilated cardiomyopathy signaling, ECM reorganization, collagen degradation and wound healing were enriched in post-R. Additionally, pathways involved in gap junction signaling, cardiac hypertrophy signaling and smooth muscle and striated muscle contraction were significantly downregulated in post-R, compared to post-NR (Figure 5C-D). Post-R also showed a significant enrichment of GP6, PKA and Eicosanoid signaling and downregulation of eNOS and PKC signaling compared to post-NR (Figure S5A-B). Increased IL4 and IL13 signaling and cardiac repair signaling pathways were observed in post-R (Figure S5C-D). Analysis of differentially methylated regions identified 295 hypomethylated and 1116 hypermethylated regions. Specifically, hypomethylation of CDH3, OPLAH and CHADL was observed, genes potentially involved in suppressing fibrillogenesis and improve cardiac health. By contrast, genes involved in microtubule and actin cytoskeleton organization and inflammation (DRAXIN, PALLD, MPO, EPSIN, etc.) were hypermethylated (Figure S5E). MAPK signaling and pathways associated with cell proliferation and differentiation were differentially hypermethylated in post-R. Pathways related to tubulin folding and microtubule-dependent protein trafficking were enriched in post-R compared to post-NR (Figure 5E-F). Examining genes that displayed concordance between expression and methylation, we observed a significant downregulation (and hypermethylation) of genes involved in microtubule and cytoskeleton signaling (MYL12A, TPM3, KIF5B), cell division (PPP1CB) and metabolism (PINK1, VPS13A) in post-R (Figure 5G). Comparing the TEM data at the post-LVAD timepoint, revealed greater sarcomere distance (Z-Z distance) in post-NR (suggestive of a damaged sarcomere), while the sarcomeric ultrastructural was preserved in post-R (Figure 5H). Gene set score computation was performed on specific sets of genes involved in actin cytoskeleton signaling and fibrotic signaling across all groups. We observed that post-R showed significant downregulation of genes associated with actin cytoskeleton signaling and fibrotic signaling (Figure 5I). Concomitantly, we also saw a significant reduction in the protein levels of TPPP3 (tubulin polymerization protein) in post-NR which remained unaffected in post-R compared to paired pre-LVAD samples. We also observed a trend towards downregulation of ECM protein, collagen1A1 in post-R compared to post-NR (Figure 5J-K). Overall, our findings suggests that reduced fibrosis and improved cytoskeletal signaling pathways play a role in promoting myocardial recovery following LVAD support.

**Figure 5.**
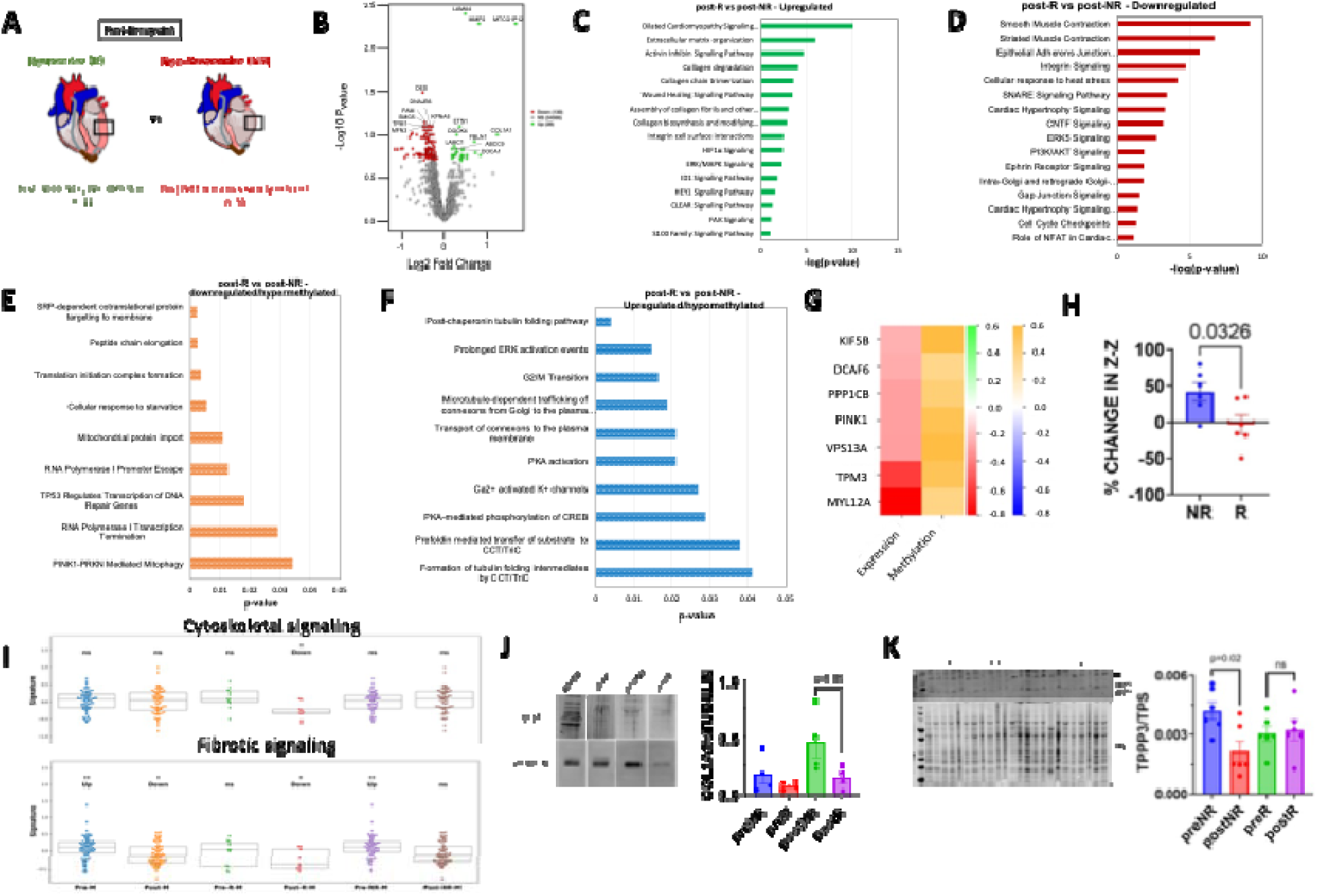
**A** Schematic of workflow for post-R (n=11) vs post-NR (n=56) comparison, **B.** Volcano plot of RNA sequencing data showing differentially expressed genes between post-R vs post-NR, **C-D.** Ingenuity pathway analysis of RNA sequencing data showing enriched pathways among upregulated and downregulated genes, **E-F.** Reactome pathway analysis showing enriched pathways among hypermethylated and hypomethylated genomic regions, **G.** Heatmap of RNA sequencing and WGBS data showing genes with concordant changes, where increased expression is associated with hypomethylation and decreased expression with hypermethylation, **H.** Quantification of percentage change in distance between the Z-lines (n=6 each), **I.** Gene set enrichment data for genes involved in cytoskeletal and fibrotic signaling, **J.** Representative western blot image and quantification of collagen1a (n=6 each), **K.** Representative western blot image and quantification of TPPP3 (n=6 each). p-value: unpaired t-test/One-way ANOVA.

### Cardiac fibrosis and microtubule remodeling pathways contribute to recovery in a mouse cardiac recovery model

We used a mouse model of cardiac recovery to investigate a conserved gene expression signature of recovery. To recapitulate the human HF cohort, we induced HF in mice by transverse aortic constriction (TAC), defined as LVEF<35%. Whereas the subset of these HF mice underwent removal of the constriction (de-TAC) served as the myocardial recovery group (LVEF>40%) (Figure 6A-B). Differential expression analysis revealed significant upregulation of 560 genes and downregulation of 497 genes in de-TAC hearts, compared to TAC hearts. Genes involved in metabolism (*Fmo5, Pon3*), cytokine signaling (*Trim34b*) and mitochondrial biogenesis (*Atp6, Nd4*) were significantly upregulated in de-TAC mice compared to TAC mice. Additionally, genes involved in interferon-gamma signaling (*Trim6, Trim34)* and cell cycle regulation (*Melk, Nudc)* were significantly downregulated in the de-TAC cohort (Figure 6C). Oxidative phosphorylation, WNT/β-catenin, sirtuin signaling, ILK signaling, mitochondrial biogenesis, NOTCH signaling and fatty acid metabolism pathways were enriched in de-TAC mice, similar to what was observed in post-R. Similarly, fibrotic signaling, integrin signaling, apoptosis, ECM remodeling and glycolysis were significantly downregulated in de-TAC mice (Figure 6D-E, Figure S6A-D). RNA sequencing data also showed a significant reduction in collagen (*Col4a1*) and Masson’s trichrome staining on de-TAC mice showed a trend towards reduced myocardial fibrosis compared to TAC mice (Figure S6E), thereby reiterating, reduced interstitial fibrotic signaling and improved cytoskeletal remodeling signaling as mechanisms implicated in cardiac recovery.

**Figure 6.**
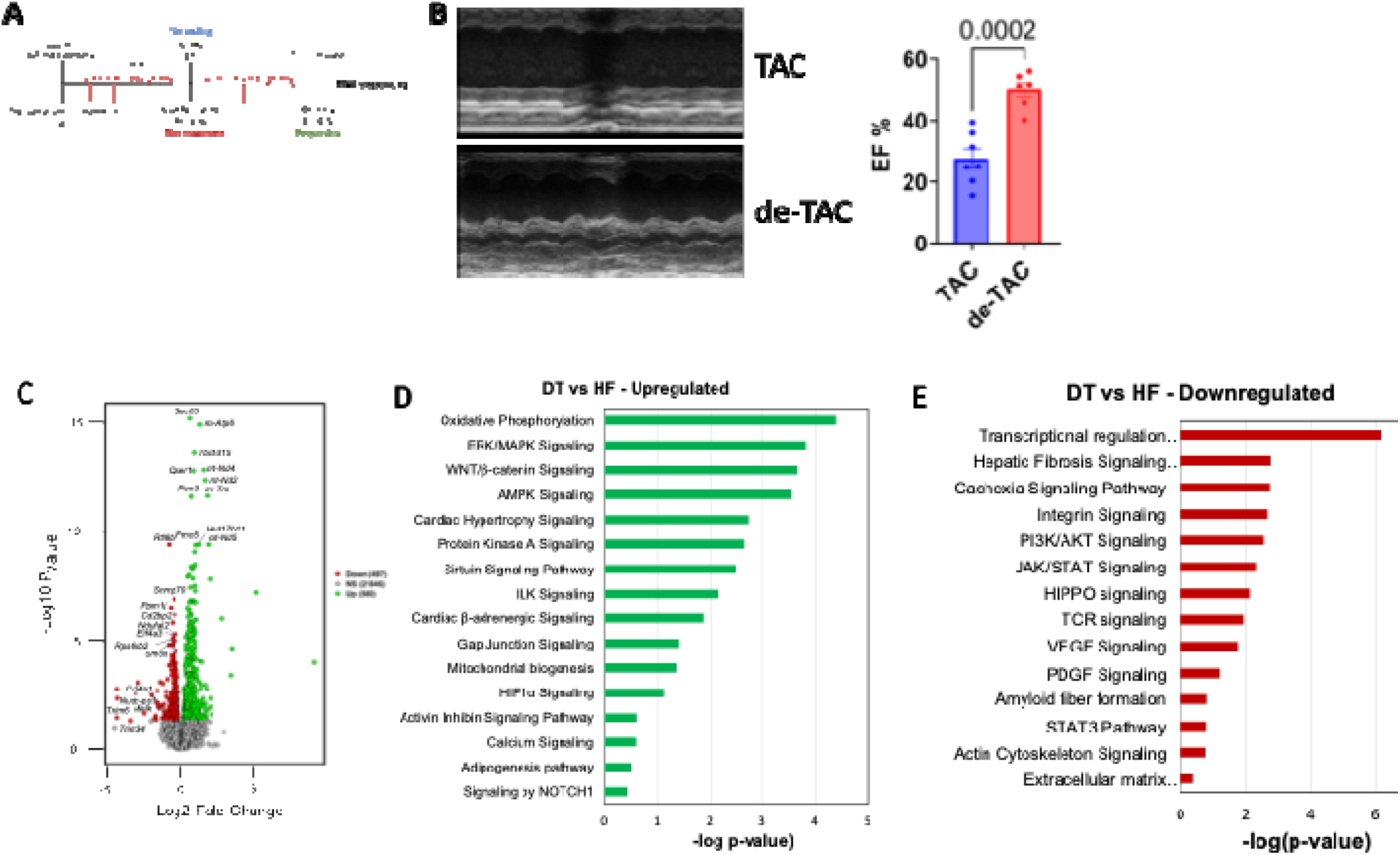
**A** Schematic of TAC (HF) and de-TAC (DT) workflow, **B.** Representative m-mode image and ejection fraction (EF%) data showing difference in cardiac structure and function (n=7 TAC, n=6 de-TAC), **C.** Volcano plot of RNA sequencing data showing differentially expressed genes between de-TAC and TAC samples, **D-E.** Ingenuity pathway analysis of RNA sequencing data showing enriched pathways among upregulated and downregulated genes.

## Discussion

DNA methylation is a key epigenetic modification that has gained attention as a diagnostic, prognostic and predictive marker in clinical research, primarily in oncology^53^. DNA methylation changes have been well-studied in various human cardiac diseases including coronary heart disease, myocardial infarction, hypertension, and HF^54,55,56^. Epigenetic alterations serve as promising biomarkers due to their stability, detectability in circulation, and potential reversibility^57^. Several predictive models incorporating DNA methylation changes have been used to assess clinical risk in CVD and HF^57^. Yet, little is known about how DNA methylation distinguishes responders from non-responders after LVAD therapy. Understanding DNA modifications observed in responders to LVAD therapy reveals epigenetic targets that may be therapeutically leveraged to promote myocardial recovery in broader HF patient populations. Therefore, in this study, we investigated DNA methylation changes, prior to and following LVAD support and integrated paired transcriptomic data to gain further insights into the molecular mechanisms underlying myocardial recovery.

Overall, our data suggests that ECM reverse remodeling and actin cytoskeletal reorganization are the key pathways observed in patients that respond favorably to LVAD therapy. Further comparison of differentially regulated pathways between post-R and post-NR samples, suggested that gap junction signaling and hypertrophy signaling were downregulated in post-R compared to post-NR. By contrast, genes related to tubulin folding intermediates and microtubule-dependent connexon trafficking, which are crucial for electrical and metabolic coupling of the actin-tubulin and contractile network^58,59,60^, were hypomethylated. Western blot data showed a reduction in collagen in post-R compared to post-NR and increased TPPP3 expression in post-NR, further validates our hypothesis of reduced fibrosis and improved microtubule architecture as key contributors to beneficial reverse cardiac remodeling.

We also observed potentially beneficial DNA methylation and transcriptomic changes in HF patients following LVAD therapy, independent of their cardiac functional response to this therapy (i.e. responders or non-responders). In all comers, RNA sequencing data showed upregulation of Hippo signaling, Notch signaling, fatty acid oxidation, cardioprotective inflammatory signaling (IL13 and IL22), myogenesis and inhibition of specific MMPs following LVAD therapy compared to pre-LVAD timepoint, potentially indicating ECM degradation and cardiac reverse remodeling pathways as effects of mechanical unloading. Additionally, downregulation of syndecan signaling, amyloid fiber formation, ceramide and glucose metabolism suggests a shift towards improved cardiac function post-LVAD therapy. WGBS data further corroborated these observations, demonstrating activation of cardioprotective Notch signaling^61^ and suppression of apoptosis and integrin pathways. Our histological findings showed a significant reduction in myocardial interstitial fibrosis and a preserved microstructural architecture in advanced HF patients following LVAD therapy.

To understand the specific molecular changes that may contribute to pathological remodeling in the non-responder population, we compared pre-NR to post-NR (paired samples). RNA sequencing data of NRs to LVAD therapy, showed a significant activation of fatty acid oxidation, apoptosis, MAPK and VEGF signaling, along with downregulation of glycolysis, gap junction and actin cytoskeleton signaling suggesting these pathways as common response to LVAD unloading. WGBS data revealed upregulation of pathways related to fetal reprograming, including mesoderm and neural plate border formation^62^ in post-NR. Furthermore, hypermethylation of genes associated with microtubule bundling and apoptosis was observed. Despite no noticeable differences in myocardial interstitial fibrosis, TEM data revealed significant alterations in sarcomere length (Z-Z distance), indicating a stretched myocardium suggestive of impaired contractile efficiency.

Conversely, in cardiac functional responders to LVAD therapy, RNA sequencing data showed enrichment in Hippo signaling, myogenesis and collagen degradation pathways, alongside suppression of DCM signaling, integrin and NFAT signaling highlighting a unique fibrosis reversal/ECM reverse remodeling and cardioprotective signaling mechanisms. While glycolytic and ceramide signaling were downregulated, glycosaminoglycan and heme signaling were activated in post-R compared to their pre-LVAD timepoint. WGBS data further showed hypermethylation of genes involved in glucose metabolism, unstable tubulin folding and microtubule-dependent trafficking of gap junction proteins. By contrast, the classic *PKM*^63^ and *KL*^64^ genes were hypomethylated (or activated) suggesting cardio protection via anti-apoptotic and increased energy production mechanisms. Histological data showed reduced myocardial fibrosis in post-R compared to pre-R.

To understand if responders were potentially predisposed to cardiac recovery and remission of HF following LVAD therapy, we then compared transcriptional and epigenetic changes between pre-R and pre-NR. Differential gene expression analysis revealed significant upregulation of PPAR signaling and anti-inflammatory signaling in pre-R. Furthermore, fibrotic signaling, pathological remodeling and hypertrophy signaling pathways were downregulated in pre-R. Genes involved in apoptotic signaling were hypermethylated, while FOXO signaling genes, previously shown to be cardioprotective under stress conditions^65,66^, were hypomethylated in pre-R. No differences in myocardial fibrosis levels and sarcomere length were observed between these two groups suggesting despite structural similarity, the responders have unique cardioprotective gene signatures that may have predisposed them to myocardial recovery following LVAD therapy.

Our findings in human HF were further supported by our HF and recovery animal model, where animals that exhibited cardiac functional recovery also showed downregulation of genes involved in fibrosis, autophagy and impaired tissue contractility (*Cryab*^67,68^*, Cltb*^69^*, Calm3*^70^). We also observed a trend towards reduced myocardial interstitial fibrosis in mice that showed recovery from HF, reinforcing the importance of cytoskeletal reorganization and fibrosis reversal for myocardial recovery.

Overall, our findings from WGBS and transcriptomic studies suggest that reduced fibrotic signaling and enhanced actin-cytoskeleton remodeling may contribute to myocardial recovery following LVAD therapy. While LVAD therapy induces a spectrum of beneficial cardioprotective and structural remodeling in advanced HF patients, only a subset of these patients achieves significant structural and functional recovery. Our data indicates unique DNA methylation patterns that could impair/drive myocardial recovery in HF patients. Specifically, non-responders showed activation of pathways that leads of increased myocardial fibrosis and a stretched myocardium, whereas responders to LVAD therapy showed genetic and epigenetic signatures of both well-organized ECM and preserved myofilament function, highlighting these two pathways as involved in reverse remodeling and myocardial recovery (Figure S7). Future studies targeting epigenetic modifications that help in fibrosis reduction and cytoskeletal remodeling may help develop novel therapeutic targets to enhance myocardial recovery rates in broader HF patient populations.

## Supplemental figures and figure legends

**Supplemental Figure 1:**
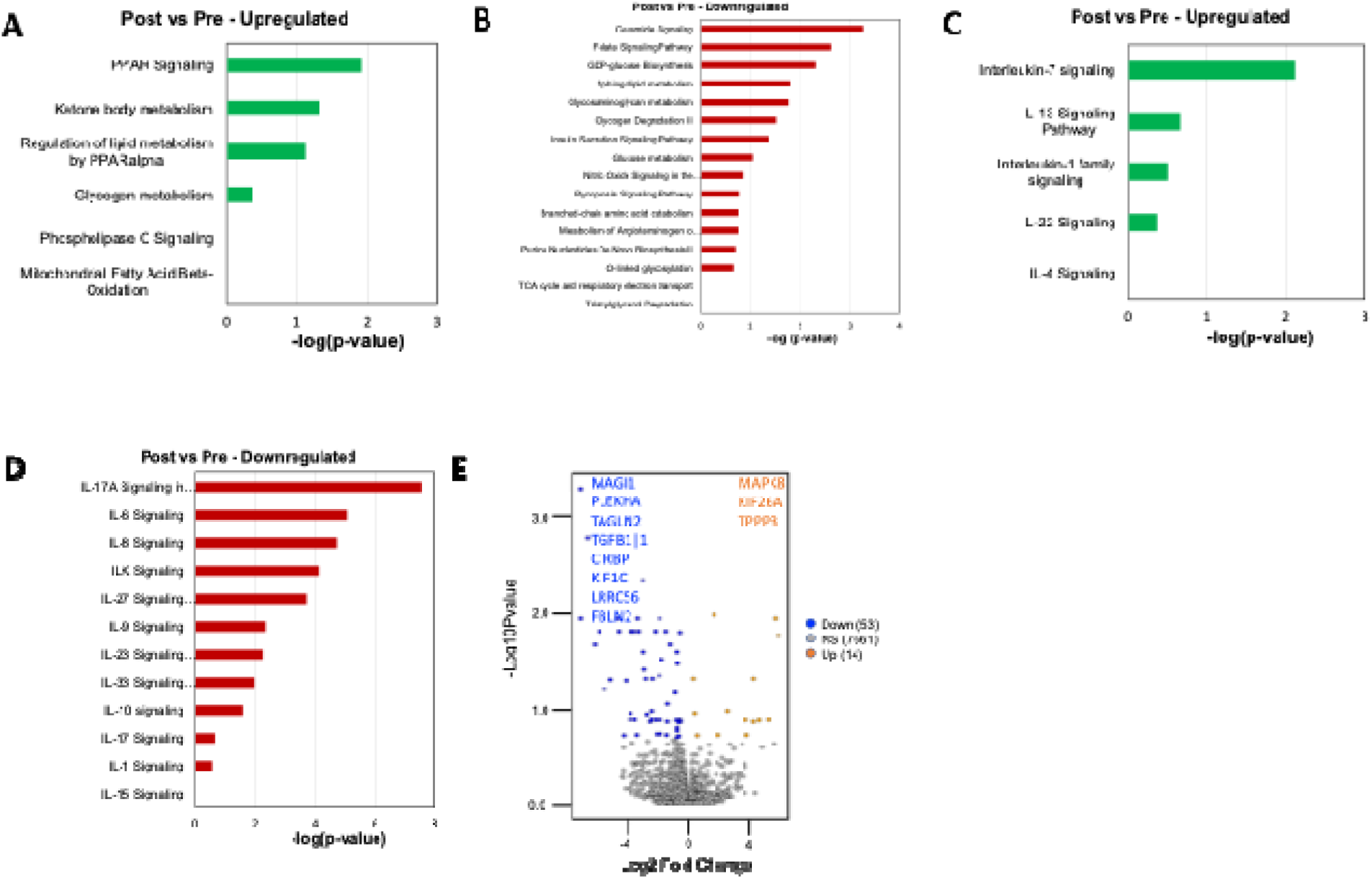
**A-D** Ingenuity pathway analysis of RNA sequencing data showing enriched pathways among upregulated and downregulated genes, in post-LVAD vs pre-LVAD patients, **E.** Volcano plot showing differentially methylated regions in post-LVAD vs pre-LVAD comparison.

**Supplemental Figure 2:**
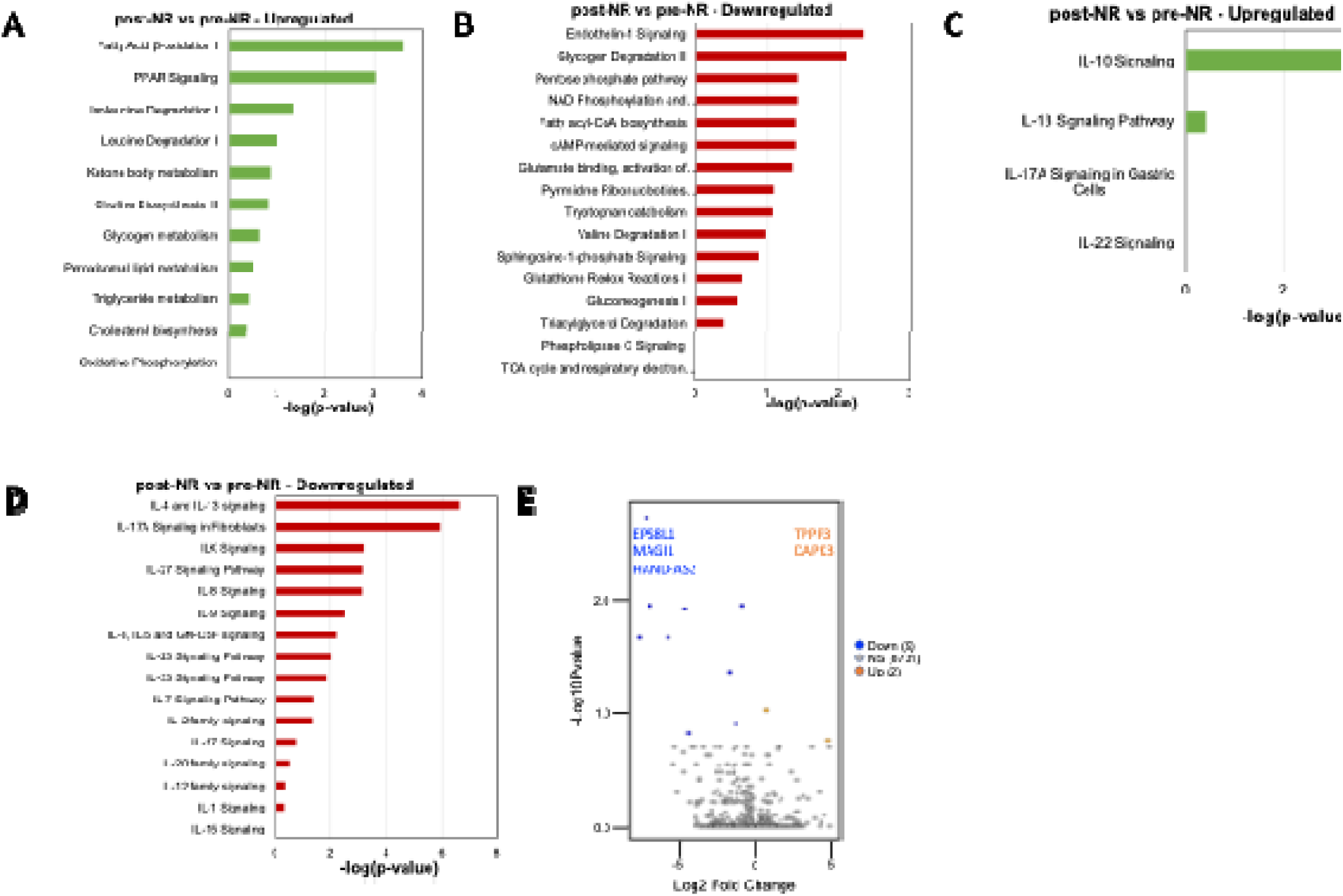
**A-D** Ingenuity pathway analysis of RNA sequencing data showing enriched pathways among upregulated and downregulated genes, in post-NR vs pre-NR, **E.** Volcano plot showing differentially methylated regions in post-NR vs pre-NR comparison.

**Supplemental Figure 3:**
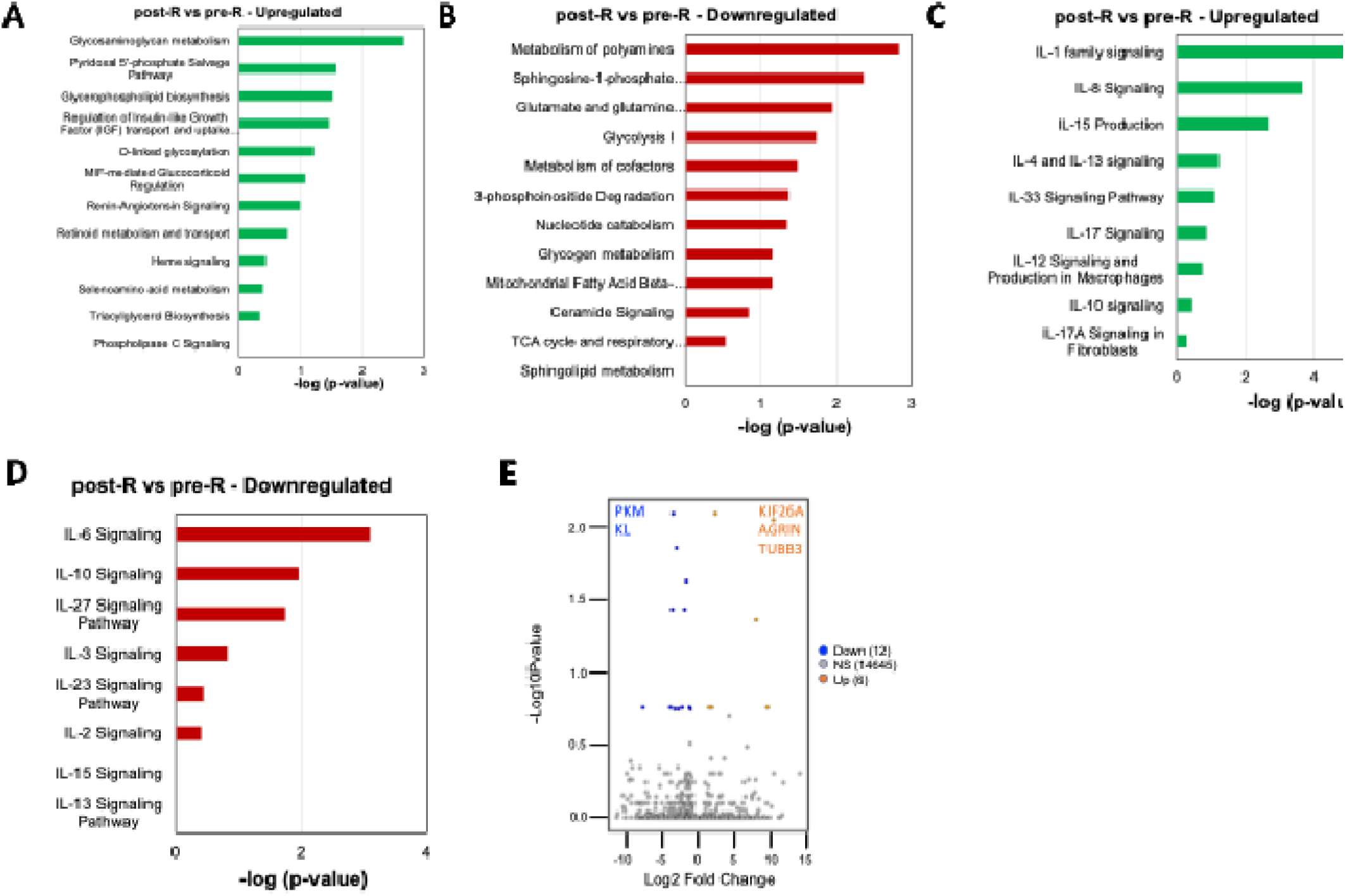
**A-D** Ingenuity pathway analysis of RNA sequencing data showing enriched pathways among upregulated and downregulated genes, in post-R vs pre-R, **E.** Volcano plot showing differentially methylated regions in post-R vs pre-R comparison.

**Supplemental Figure 4:**
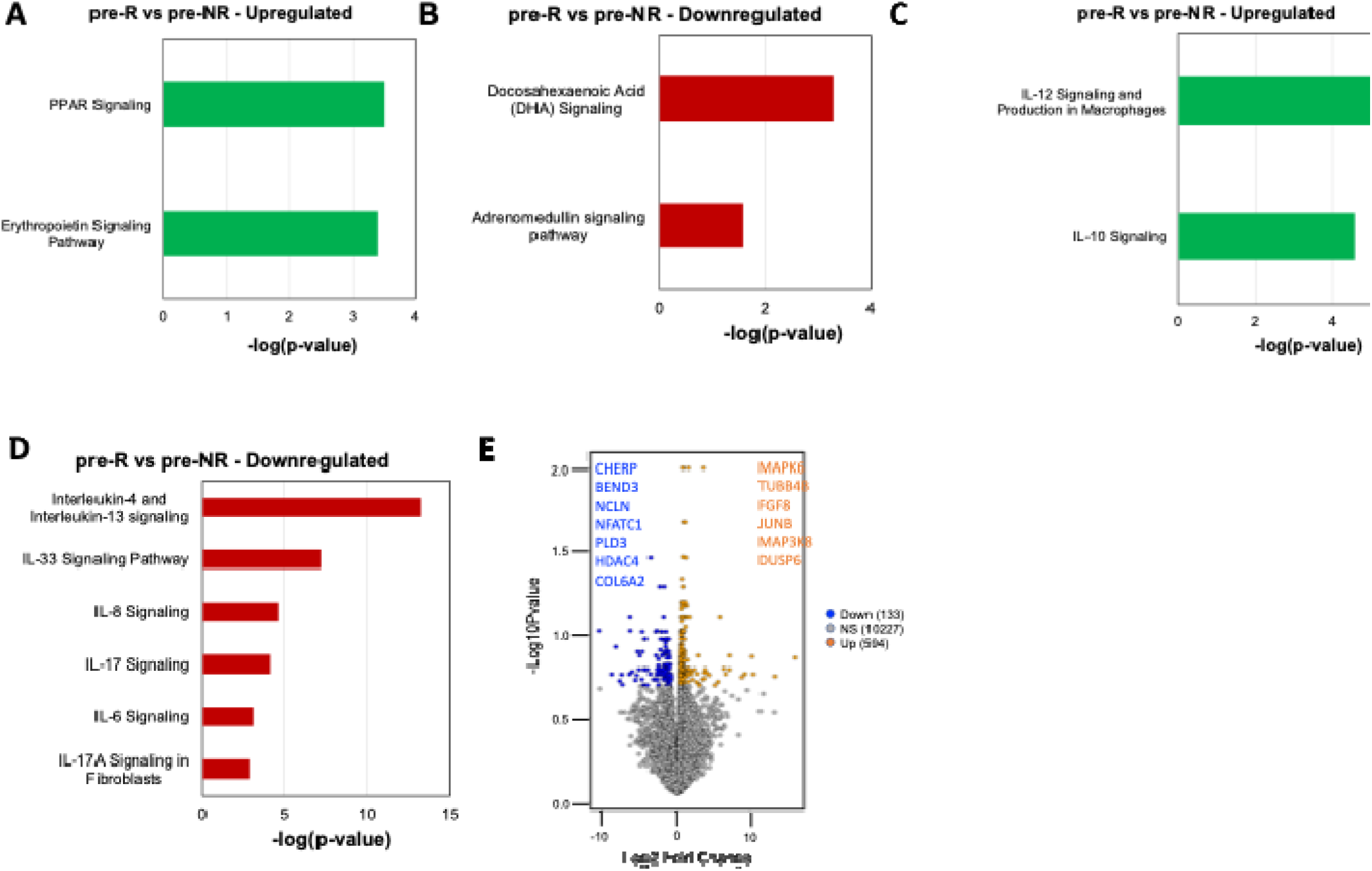
**A-D** Ingenuity pathway analysis of RNA sequencing data showing enriched pathways among upregulated and downregulated genes, in pre-R vs pre-NR, **E.** Volcano plot showing differentially methylated regions in pre-R vs pre-NR comparison.

**Supplemental Figure 5:**
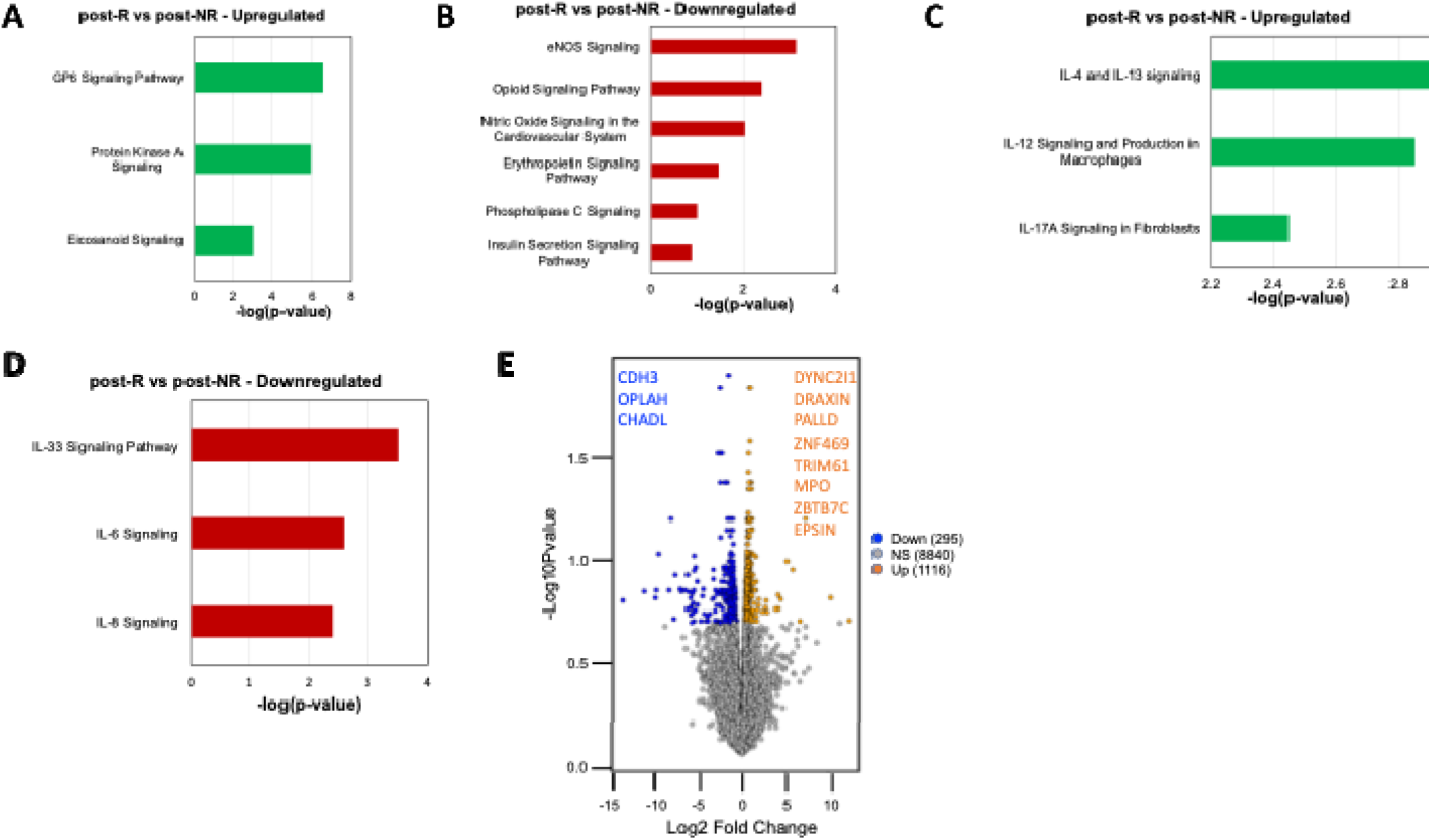
**A-D** Ingenuity pathway analysis of RNA sequencing data showing enriched pathways among upregulated and downregulated genes, in post-R vs post-NR, **E.** Volcano plot showing differentially methylated regions in post-R vs post-NR comparison.

**Supplemental Figure 6:**
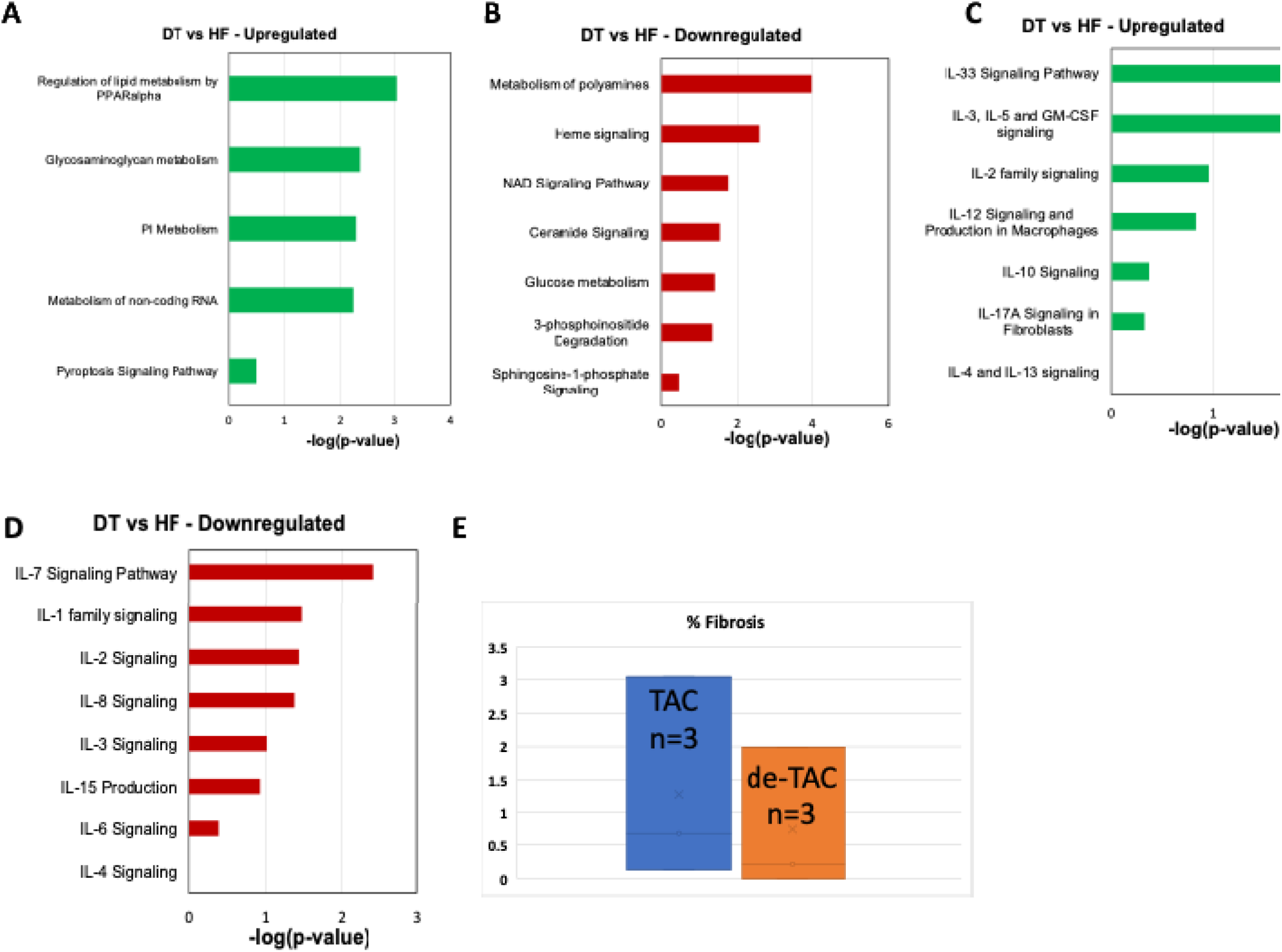
**A-D** Ingenuity pathway analysis of RNA sequencing data showing enriched pathways among upregulated and downregulated genes, in de-TAC vs TAC, **E.** Masson’s trichrome quantification data between de-TAC and TAC samples (n=3 each). p-value: unpaired t-test.

**Supplemental Figure 7:**
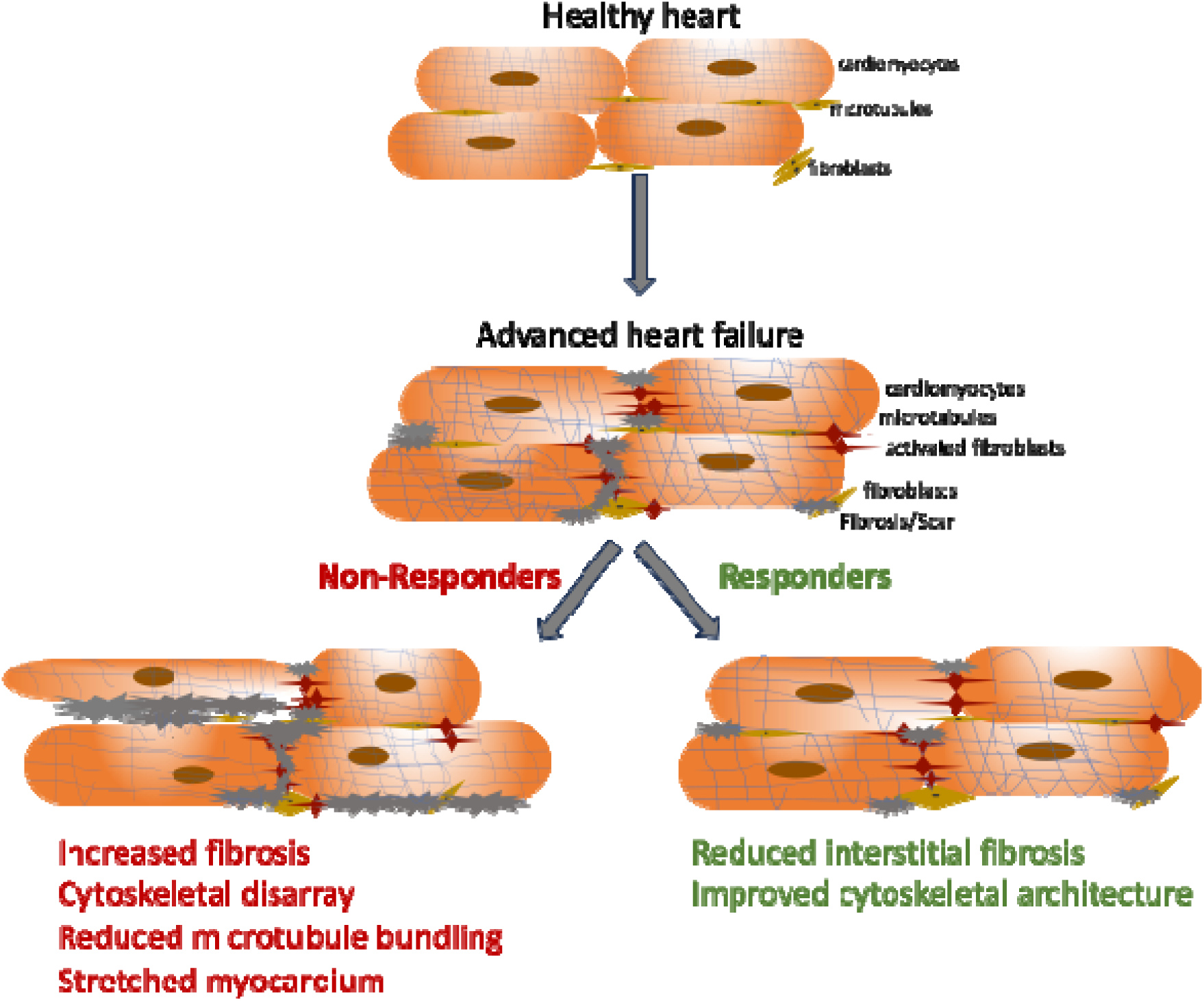
Increased fibrosis and disrupted cytoskeletal signaling maybe potential barriers to myocardial recovery following LVAD therapy.

## Acknowledgements

We thank the Nora Eccles Harrison Cardiovascular Research and Training Institute and the Nora Eccles Harrison Treadwell Foundation for their support of this research project. We acknowledge funding from the following sources: American Heart Association (AHA) postdoctoral fellowship 23POST1019351 awarded to T.S.S., National Institutes of Health (NIH) R01HL135121 and R01HL132067 to SGD.

## References

1. Yan, T., et al. Burden, Trends, and Inequalities of Heart Failure Globally, 1990 to 2019: A Secondary Analysis Based on the Global Burden of Disease 2019 Study. J. Am. Heart Assoc. 12, e027852 (2023).

2. Shahim, B., Kapelios, C. J., Savarese, G. & Lund, L. H. Global Public Health Burden of Heart Failure: An Updated Review. Card. Fail. Rev. 9, e11 (2023).

3. Yang, Y., et al. The Present Clinical Treatment and Future Emerging Interdisciplinary for Heart Failure: Where we are and What we can do. Intensive Care Res. 3, 3–11 (2023).

4. Shah, P., et al. Framework to Classify Reverse Cardiac Remodeling With Mechanical Circulatory Support: The Utah-Inova Stages. Circ. Heart Fail. 14, e007991 (2021).

5. Kyriakopoulos, C. P. et al. Predicting Cardiac Structural And Functional Improvement On Left Ventricular Assist Device Support: The Externally Validated UCAR Score. J. Card. Fail. 28, S55 (2022).

6. Wever-Pinzon, O., et al. Cardiac Recovery During Long-Term Left Ventricular Assist Device Support. J. Am. Coll. Cardiol. 68, 1540–1553 (2016).

7. Topkara, V. K., et al. Myocardial Recovery in Patients Receiving Contemporary Left Ventricular Assist Devices: Results from the Interagency Registry for Mechanically Assisted Circulatory Support (INTERMACS). Circ. Heart Fail. 9, 10.1161/CIRCHEARTFAILURE.116.003157 e003157 (2016).

8. Marinescu, K. K., Uriel, N., Mann, D. L. & Burkhoff, D. Left ventricular assist device-induced reverse remodeling: it’s not just about myocardial recovery. Expert Rev. Med. Devices 14, 15–26 (2017).

9. Taleb, I., Tseliou, E., Fang, J. C. & Drakos, S. G. A Mechanical Bridge to Recovery as a Bridge to Discovery: Learning From Few and Applying to Many. Circulation 145, 562–564 (2022).

10. Drakos, S. G., Pagani, F. D., Lundberg, M. S. & Baldwin, J. T. Advancing the Science of Myocardial Recovery With Mechanical Circulatory Support: A Working Group of the National, Heart, Lung, and Blood Institute. JACC Basic Transl. Sci. 2, 335–340 (2017).

11. Wei, Y., Cao, H., Peng, Y.-Y. & Zhang, B. Alterated gene expression in dilated cardiomyopathy after left ventricular assist device support by bioinformatics analysis. Front. Cardiovasc. Med. 10, 1013057 (2023).

12. Birks, E. J. Molecular Changes After Left Ventricular Assist Device Support for Heart Failure. Circ. Res. 113, 777–791 (2013).

13. Willmer, T., Mabasa, L., Sharma, J., Muller, C. J. F. & Johnson, R. Blood-Based DNA Methylation Biomarkers to Identify Risk and Progression of Cardiovascular Disease. Int. J. Mol. Sci. 26, 2355 (2025).

14. Agha, G., et al. Blood Leukocyte DNA Methylation Predicts Risk of Future Myocardial Infarction and Coronary Heart Disease. Circulation 140, 645–657 (2019).

15. Pepin, M. E., et al. DNA methylation reprograms cardiac metabolic gene expression in end-stage human heart failure. Am. J. Physiol.-Heart Circ. Physiol. 317, H674–H684 (2019).

16. Liao, X., et al. Effect of mechanical unloading on genome-wide DNA methylation profile of the failing human heart. JCI Insight 8, e161788.

17. Movassagh, M., et al. Differential DNA Methylation Correlates with Differential Expression of Angiogenic Factors in Human Heart Failure. PLOS ONE 5, e8564 (2010).

18. Haas, J., et al. Alterations in cardiac DNA methylation in human dilated cardiomyopathy. EMBO Mol. Med. 5, 413–429 (2013).

19. Liu, C.-F. & Tang, W. H. W. Epigenetics in Cardiac Hypertrophy and Heart Failure. JACC Basic Transl. Sci. 4, 976–993 (2019).

20. Kanwar, M. K., et al. Clinical myocardial recovery in advanced heart failure with long term left ventricular assist device support. J. Heart Lung Transplant. 41, 1324–1334 (2022).

21. Topkara, V. K., et al. Machine Learning-Based Prediction of Myocardial Recovery in Patients With Left Ventricular Assist Device Support. Circ. Heart Fail. 15, e008711 (2022).

22. Diakos, N. A., et al. Evidence of Glycolysis Up-Regulation and Pyruvate Mitochondrial Oxidation Mismatch During Mechanical Unloading of the Failing Human Heart: Implications for Cardiac Reloading and Conditioning. JACC Basic Transl. Sci. 1, 432–444 (2016).

23. Dobin, A., et al. STAR: ultrafast universal RNA-seq aligner. Bioinformatics 29, 15–21 (2013).

24. Leek, J. T., Johnson, W. E., Parker, H. S., Jaffe, A. E. & Storey, J. D. The sva package for removing batch effects and other unwanted variation in high-throughput experiments. Bioinformatics 28, 882–883 (2012).

25. Love, M. I., Huber, W. & Anders, S. Moderated estimation of fold change and dispersion for RNA-seq data with DESeq2. Genome Biol. 15, 550 (2014).

26. Reiner, A., Yekutieli, D. & Benjamini, Y. Identifying differentially expressed genes using false discovery rate controlling procedures. Bioinformatics 19, 368–375 (2003).

27. Krämer, A., Green, J., Pollard, J. & Tugendreich, S. Causal analysis approaches in Ingenuity Pathway Analysis. Bioinforma. Oxf. Engl. 30, 523–530 (2014).

28. Krueger, F. & Andrews, S. R. Bismark: a flexible aligner and methylation caller for Bisulfite-Seq applications. Bioinformatics 27, 1571–1572 (2011).

29. Akalin, A., et al. methylKit: a comprehensive R package for the analysis of genome-wide DNA methylation profiles. Genome Biol. 13, R87 (2012).

30. Shankar, T. S., et al. Cardiac-specific deletion of voltage dependent anion channel 2 leads to dilated cardiomyopathy by altering calcium homeostasis. Nat. Commun. 12, 4583 (2021).

31. Visker, J. R., et al. Enhancing mitochondrial pyruvate metabolism ameliorates ischemic reperfusion injury in the heart. JCI Insight 9, e180906 (2024).

32. Wever-Pinzon, O., et al. Magnitude and Time Course of Changes Induced by Continuous-Flow Left Ventricular Assist Device Unloading in Chronic Heart Failure: Insights into Cardiac Recovery. J. Heart Lung Transplant. 32, S31–S32 (2013).

33. Shankar, T. S., et al. Cardiac-specific deletion of voltage dependent anion channel 2 leads to dilated cardiomyopathy by altering calcium homeostasis. Nat. Commun. 12, 4583 (2021).

34. Drakos, S. G., et al. Impact of mechanical unloading on microvasculature and associated central remodeling features of the failing human heart. J. Am. Coll. Cardiol. 56, 382–391 (2010).

35. Miao, C., et al. Pro- and anti-fibrotic effects of vascular endothelial growth factor in chronic kidney diseases. Ren. Fail. 44, 881–892.

36. Bakalenko, N., Kuznetsova, E. & Malashicheva, A. The Complex Interplay of TGF-β and Notch Signaling in the Pathogenesis of Fibrosis. Int. J. Mol. Sci. 25, 10803 (2024).

37. Alvarez-Argote, S., et al. IL-13 promotes functional recovery after myocardial infarction via direct signaling to macrophages. JCI Insight 9, e172702 (2024).

38. Che, Y., et al. IL-22 ameliorated cardiomyocyte apoptosis in cardiac ischemia/reperfusion injury by blocking mitochondrial membrane potential decrease, inhibiting ROS and cytochrome C. Biochim. Biophys. Acta BBA - Mol. Basis Dis. 1867, 166171 (2021).

39. Zhou, X., Wan, L., Xu, Q., Zhao, Y. & Liu, J. Notch signaling activation contributes to cardioprotection provided by ischemic preconditioning and postconditioning. J. Transl. Med. 11, 251 (2013).

40. Badolia, R., et al. The Role of Nonglycolytic Glucose Metabolism in Myocardial Recovery Upon Mechanical Unloading and Circulatory Support in Chronic Heart Failure. Circulation 142, 259–274 (2020).

41. Borodzicz-Jażdżyk, S., Jażdżyk, P., Łysik, W., Cudnoch-J drzejewska, A. & Czarzasta, K. Sphingolipid metabolism and signaling in cardiovascular diseases. Front. Cardiovasc. Med. 9, 915961 (2022).

42. Oláh, J., Lehotzky, A., Szénási, T., Berki, T. & Ovádi, J. Modulatory Role of TPPP3 in Microtubule Organization and Its Impact on Alpha-Synuclein Pathology. Cells 11, 3025 (2022).

43. Caporizzo, M. A. & Prosser, B. L. The microtubule cytoskeleton in cardiac mechanics and heart failure. Nat. Rev. Cardiol. 19, 364–378 (2022).

44. Abe, J., et al. MAGI1 as a link between endothelial activation and ER stress drives atherosclerosis. https://insight.jci.org/articles/view/125570/pdf (2019) doi:10.1172/jci.insight.125570.

45. Muscle Contraction and Locomotion | Biology for Majors II. https://courses.lumenlearning.com/suny-wmopen-biology2/chapter/muscle-contraction-and-locomotion/.

46. Increased regulatory activity of the calcineurin/NFAT pathway in human heart failure - Diedrichs - 2004 - European Journal of Heart Failure - Wiley Online Library. https://onlinelibrary.wiley.com/doi/10.1016/j.ejheart.2003.07.007.

47. Lugrin, J., et al. The systemic deletion of interleukin-1α reduces myocardial inflammation and attenuates ventricular remodeling in murine myocardial infarction. Sci. Rep. 13, 4006 (2023).

48. Ameri, K., et al. Administration of Interleukin-15 Peptide Improves Cardiac Function in a Mouse Model of Myocardial Infarction. J. Cardiovasc. Pharmacol. 75, 98–102 (2020).

49. Olejnik, A., Franczak, A., Krzywonos-Zawadzka, A., Kałużna-Oleksy, M. & Bil-Lula, I. The Biological Role of Klotho Protein in the Development of Cardiovascular Diseases. BioMed Res. Int. 2018, 5171945 (2018).

50. Li, Q., et al. PKM1 Exerts Critical Roles in Cardiac Remodeling under Pressure Overload in the Heart. Circulation 144, 712–727 (2021).

51. Foxo Transcription Factors Blunt Cardiac Hypertrophy by Inhibiting Calcineurin Signaling | Circulation. https://www.ahajournals.org/doi/10.1161/circulationaha.106.637124.

52. Abeyrathna, P. & Su, Y. The Critical Role of Akt in Cardiovascular Function. Vascul. Pharmacol. 74, 38–48 (2015).

53. Galbraith, K. & Snuderl, M. DNA methylation as a diagnostic tool. Acta Neuropathol. Commun. 10, 71 (2022).

54. Guarrera, S., et al. Gene-specific DNA methylation profiles and LINE-1 hypomethylation are associated with myocardial infarction risk. Clin. Epigenetics 7, 133 (2015).

55. Westerman, K., et al. DNA methylation modules associate with incident cardiovascular disease and cumulative risk factor exposure. Clin. Epigenetics 11, 142 (2019).

56. Krolevets, M., et al. DNA methylation and cardiovascular disease in humans: a systematic review and database of known CpG methylation sites. Clin. Epigenetics 15, 56 (2023).

57. Desiderio, A., et al. DNA methylation in cardiovascular disease and heart failure: novel prediction models? Clin. Epigenetics 16, 115 (2024).

58. Marchal, G. A., Galjart, N., Portero, V. & Remme, C. A. Microtubule plus-end tracking proteins: novel modulators of cardiac sodium channels and arrhythmogenesis. Cardiovasc. Res. 119, 1461–1479 (2023).

59. Kelly, J. J., et al. Snapshots of actin and tubulin folding inside the TRiC chaperonin. Nat. Struct. Mol. Biol. 29, 420–429 (2022).

60. Epifantseva, I. & Shaw, R. M. Intracellular trafficking pathways of Cx43 gap junction channels. Biochim. Biophys. Acta BBA - Biomembr. 1860, 40–47 (2018).

61. Olejnik, A., Radajewska, A., Krzywonos-Zawadzka, A. & Bil-Lula, I. Klotho inhibits IGF1R/PI3K/AKT signalling pathway and protects the heart from oxidative stress during ischemia/reperfusion injury. Sci. Rep. 13, 20312 (2023).

62. Brade, T., Pane, L. S., Moretti, A., Chien, K. R. & Laugwitz, K.-L. Embryonic Heart Progenitors and Cardiogenesis. Cold Spring Harb. Perspect. Med. 3, a013847 (2013).

63. PKM1 Exerts Critical Roles in Cardiac Remodeling Under Pressure Overload in the Heart | Circulation. https://www.ahajournals.org/doi/10.1161/CIRCULATIONAHA.121.054885#.

64. Chen, K., et al. Klotho Deficiency Causes Heart Aging via Impairing the Nrf2-GR Pathway. Circ. Res. 128, 492–507 (2021).

65. Skurk, C., et al. The FOXO3a Transcription Factor Regulates Cardiac Myocyte Size Downstream of AKT Signaling*. J. Biol. Chem. 280, 20814–20823 (2005).

66. Ni, Y. G., et al. Foxo Transcription Factors Blunt Cardiac Hypertrophy by Inhibiting Calcineurin Signaling. Circulation 114, 1159–1168 (2006).

67. Wang, X., et al. αB-Crystallin Modulates Protein Aggregation of Abnormal Desmin. Circ. Res. 93, 998–1005 (2003).

68. Wang, X., et al. AlphaB-crystallin modulates protein aggregation of abnormal desmin. Circ. Res. 93, 998–1005 (2003).

69. He, L., Chen, L. & Li, L. The mechanosensitive APJ internalization via clathrin-mediated endocytosis: A new molecular mechanism of cardiac hypertrophy. Med. Hypotheses 90, 6–10 (2016).

70. Hussey, J. W., Limpitikul, W. B. & Dick, I. E. Calmodulin Mutations in Human Disease. Channels 17, 2165278.

